# Anchorage by seed mucilage prevents seed dislodgement in high surface flow: a mechanistic investigation

**DOI:** 10.1101/2021.09.25.461784

**Authors:** Vincent S. Pan, Cecilia Girvin, Eric F. LoPresti

## Abstract

1. *Background and Aims:* Seed mucilage is a common and highly diverse trait shared among thousands of angiosperm species. While long recognized that mucilage allows seeds to anchor to substrates (antitelechory), resisting abiotic and biotic dislodgement, we still lack a mechanistic understanding of this process.
2. *Methods:* We propose a mechanistic model of how mucilage affects substrate anchorage and fluid resistance, ultimately contributing to dislodgement resistance. To test this model, we subjected mucilaginous seeds of 52 species, varying in eight measured seed traits, to seven days of continuous water flow at a range of dislodgement potentials.
3. *Key Results:* Supporting our model, mucilage mass increased force necessary to dislodge both dry and wet seeds; our measurement of the dislodgement force of dry mucilage explained time to dislodgement well. The effect size was remarkably large; increasing the standardized mucilage mass by one standard deviation resulted in a 280-fold increase in the time to dislodgement. Fluid resistance was largely dependent on speed of water flow and the seed’s modified drag coefficient, but not seed traits. Neither mucilage expansion speed nor mucilage decay rate explained dislodgement potential well.
4. *Conclusions:* Our results suggest that the degree of anchorage to substrate, measured with a simple dislodgement force assay, is highly predictive of mucilaginous seed retention in highly erosive environments. In contrast, we found that other seed and mucilage traits are of lesser importance to anchorage.

## Introduction

Diaspore mucilage (myxodiaspory, hereafter: seed mucilage) is an extremely widespread trait in plants with thousands of species of flowering plants (Grubert, 1974, 1981). When wetted, mucilage absorbs water and swells, releasing a complex matrix of hydrated cellulosic, pectic, and hemicellulosic sugars (Western, 2012). This matrix may be sticky or slippery when wet, but after drying, generally anchors seeds to substrates tightly (Kreitschitz et al., 2021b, Pan et al., 2021; Figure 2C, F). Previous workers have observed its utility in anchorage to substrates (antitelochory) and noted its ubiquity among plants in highly erosive environments (Ellner & Schmida, 1981, Garcia-Fayos et al., 2013). In the only example explicitly dealing with antitelechory, Engelbrecht et al. (2014) found that seeds of *Fumana ericoides* (Cistaceae) from low erosion sites produced significantly less mucilage than those from high erosion sites; suggesting functional, possibly adaptive, variation in the trait, though another species in the same family (*Helianthemum violaceum*) did not show the same pattern.

Much excellent experimental research on the ecological functions of mucilage in relation to dispersal, germination, defense, and stress amelioration (e.g. Fuller & Hay, 1983, Kreitschitz et al., 2021a, Zhou et al., 2021) has been done, but only with one or a few species. Therefore, while we know many functional roles of mucilage in certain species, we do not know how specific aspects of mucilage contribute to these roles. As mucilage traits vary both within and across species, determining the functional roles of this variation can provide insight into the evolutionary pressures shaping these traits. When looking broadly, mucilage mass and dislodgement force measured across 53 plant species in LoPresti et al. (2019) and Pan et al. (2021) were strongly correlated among more-closely related species. However, when looking in- depth at close relatives, there is often much variation (Figure 2D-E). Comparisons of many *Plantago* species reveal a wide variety of mucilage structure across the genus (Cowley & Burton, 2021, Cowley et al. 2021) as does examining variation across populations of other widespread species or genera (e.g. Inceer et al., 2012; Villellas & Garcia, 2012, Poulain et al., 2019).

Therefore, examining functional roles of mucilage across closely and distantly related species allows insight into the contributions of specific measurable mucilage traits that differ between those species.

Rainfall, and subsequent surface flow across steep slopes, is a common type of dislodgement that is directional and may be responsible for staggering seed mortality. Han et al. (2011) found that 60-90% of seeds experimentally placed on shallow slopes were washed away in a single hour of heavy rain; similar results were reported by Aerts et al. (2006) who further found that the effect was most pronounced on bare slopes without leaf litter, although neither used any mucilaginous seeds in their studies. A single, eight-hour rainfall event totaling 44 mm in a 60-ha watershed in Laos was estimated to have washed 1.34 million seeds into a river due to surface flow, estimated to be >1% of all seeds in the area (de Rouw et al., 2018). Given that the area receives 1403 mm of precipitation annually (de Rouw et al., 2018), this sort of flow could disperse or cause mortality to a large fraction of seeds produced each year. We know of no comparable estimates for other areas, but this sort of erosion is certainly a risk to seeds in many, if not all, habitats, and could represent a powerful selective force on antitelechorous traits (Ellner & Schmida, 1981, Garcia-Fayos et al., 2013, Engelbrecht et al., 2014).

Imbibed seed mucilage binds seeds strongly as it dries to whatever it contacts, be it loose substrate, hard-packed substrate, rocks, leaves, or the mother plant itself (Pan et al., 2021). The anchorage strength of mucilage after a wetting-drying cycle is extremely strong and correlates positively with the amount of mucilage produced (Kreitschitz et al., 2021b, Pan et al., 2021).

Wet mucilage is still sticky, which may be important for zoochorous dispersal (Ryding, 2001), but the strength of attachment is considerably lower than after drying and has not been well characterized except in flax, *Linum usitatissimum* (Kreitschitz et al., 2015). In the face of surface flow, the speed at which mucilage rehydrates, switching from stronger to weaker attachment, may thus contribute to the length of time that substrate anchorage is effective. Over longer periods, the breakdown of the mucilage layer in water may also reduce the amount of mucilage or strength of mucilage attachment, lowering resistance to dislodgement. Seed mucilage may further alter the fluid resistance in complex ways. The expansion of the mucilage envelope provides a larger area upon which flowing water may exert force. However, some of the drag may be alleviated by the improvement in sphericity for more rigid mucilage envelopes or closer resemblance to a streamlined shape for more malleable mucilage envelopes in fast flowing water. In addition, any lubrication provided by the mucilage layer between water and seed could reduce the force that is exerted on the seed.

Integrating these functional hypotheses, we propose a multi-part model for the physical mechanisms underlying antitelechory in mucilaginous seeds (Figure 1, 2). Specifically, we work under the framework of the two major mechanistic processes, the ability of seeds to withstand horizontal shear force and the amount of shear force the seeds experience as a result of fluid resistance. Traits which contribute to the attachment potential are the amount of mucilage, the rate at which that mucilage expands, the rate it decays, and the seed size. Traits which contribute to drag are seed size (including imbibed mucilage), the mucilage decay and expansion rates, and seed shape. Externally, the intensity of surface flow also contributes to potential for successful anchorage.

**Figure 1.**
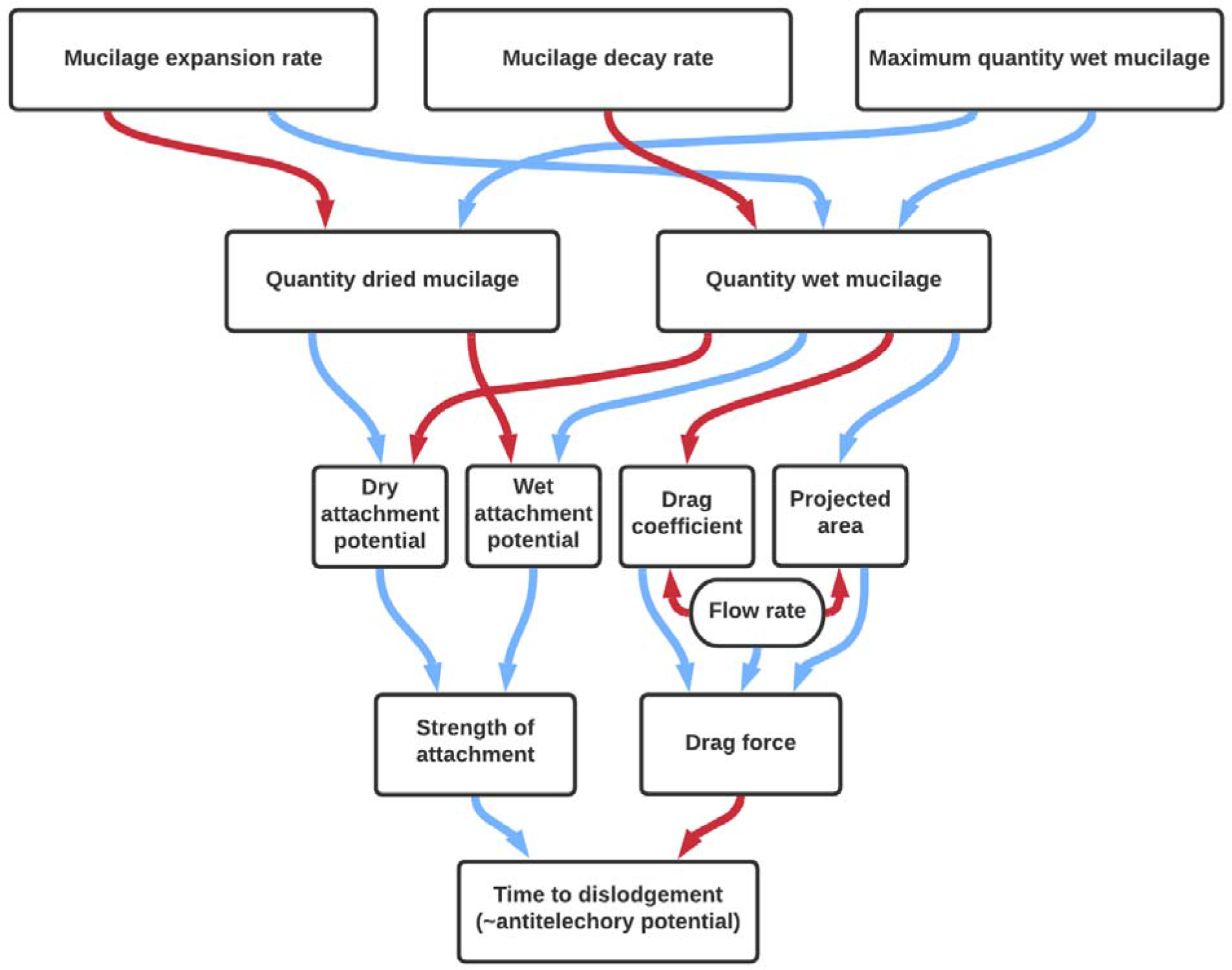
Diagram illustrating the hypothetical ways that mucilage traits may mechanistically influence seed anchorage. Red arrows represent a negative effect, whereas blue arrows represent a positive effect.

**Figure 2.**
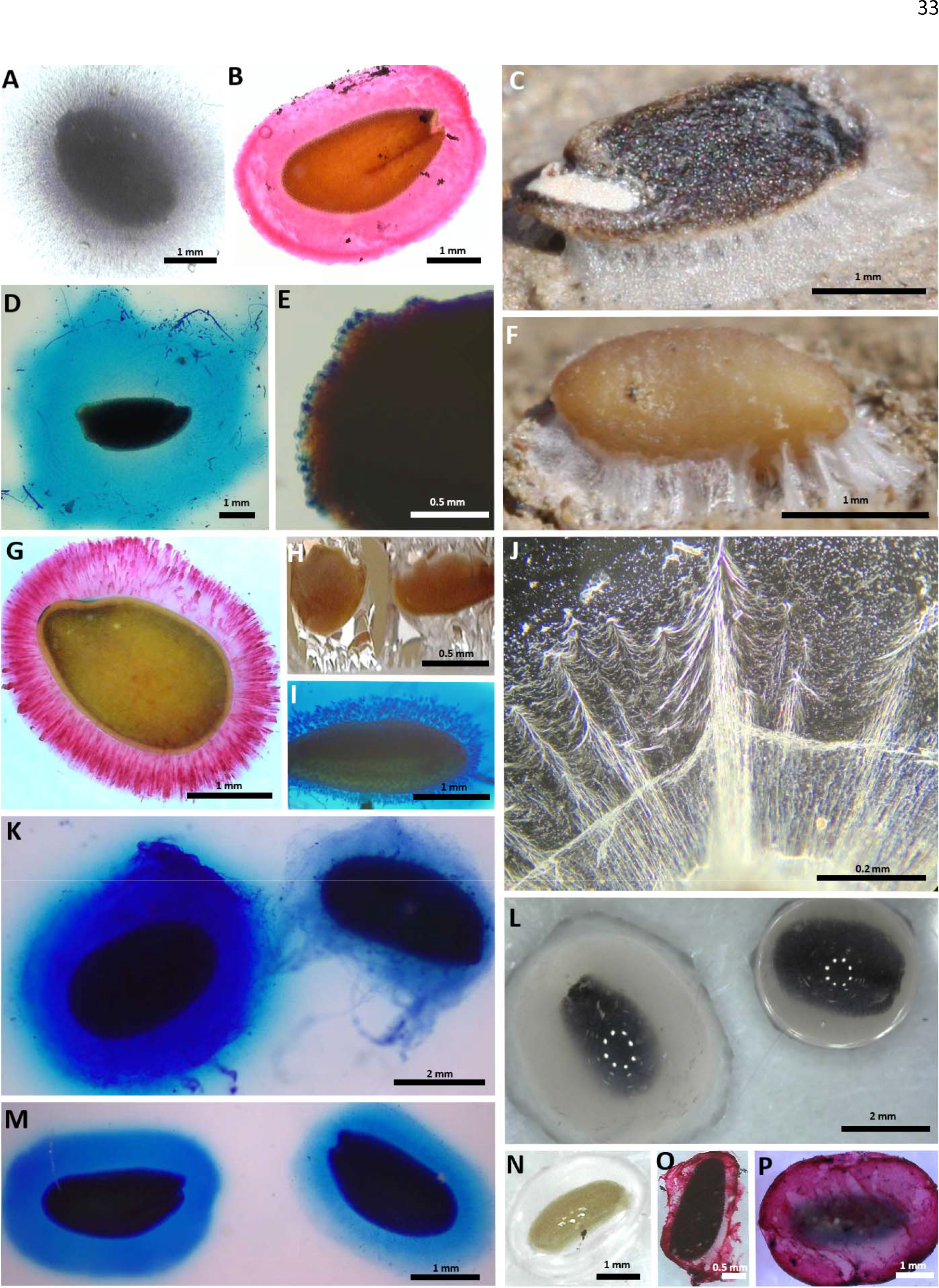
Light microscope images or photographs of (A) *Ocimum basilicum* (Lamiaceae) with visible starch granules stained with Lugol’s iodine. (B) *Lepidium sativum* (Brassicaceae) stained with Ruthenium red (RR) to show different mucilage structures based on distributions of pectic sugars. (C) *Dracocephalum moldavica* dried mucilage strongly cemented on soil surface (as part of Pan et al 2020). (D) *D. moldavica* (Lamiaceae) and (E) *D. parviflorum* both stained with methylene blue show different interspecific mucilage volume. (F) *Gilia leptantha* (Polemoniaceae) dried mucilage strongly cemented on soil surface. (G) *Linum grandiflornum* (Linaceae), RR stain, (H) *Li. grandiflorum* wet mucilage stretching apart by two pieces of filter paper. (I) *Plantago* (Plantaginaceae) seed stained with methylene blue. (J) *Le. sativum* dried mucilage magnified under phase contrast. (K) *Salvia hispanica* nutlet stained with methylene blue; Mucilage decay due to extended imbibition affects mucilage quantity: the seeds on the right that were imbibed for 12 days had a thinner mucilage envelope compared to the seeds on the left that were imbibed for an hour. Seed mucilage also varies in volume and projected area (L) two unstained *O. basilicum* nutlets from the same batch show different intraspecific mucilage volumes. (M) *Le. sativum* stained with methylene blue, same decay treatments as K. (N) *Matricaria chamomilla* (Asteraceae) achene mucilage, unstained. (O) *Melissa officinalis* (Lamiaceae), RR stain, and (P) *Salvia coccinea* (Lamiaceae), RR stain. Staining methodology adapted from (Kreitschitz et al., 2021b)

In this study, we tested the proposed physical model with mucilaginous seeds of 52 species in varying intensities of erosive surface flows. We examined how each seed trait relates to substrate attachment potential for the ability to withstand shear force and to projected area (largest cross-sectional area) and drag coefficient for fluid resistance. We characterized seed mass, seed mucilage mass, and two temporal aspects of mucilage quantity in water (i.e. speed of mucilage expansion and breakdown). After identifying useful predictors of dislodgement potential, we compared the marginal predictive utility of the functional assays to provide guidance on trait selection in future functional investigations.

## Methods

### Dislodgement resistance experiment

We experimentally tested whether seed mucilage facilitates anchorage during erosive surface flow for 52 species of mucilaginous-seeded plants from 10 families (Table S1, Figure S1). The species were selected haphazardly based on availability from commercial seed sources, our field collections, or the USDA Germplasm Resources Information Network program (USDA-ARS, 2015). Preliminary experiments in the field (see supplement) showed that seeds can remain stuck to inclined sandstone, vertical clay cliffs, and frequently flooded riverbanks during several rainstorms (Figure 3A, D-E). To mimic the erosive effect of surface flow during these rainstorm events, we developed a simple laboratory assay that subjected anchored seeds of each species to a wide range of surface flow rates that seeds may encounter in the field (Figure S2). Dry seeds were imbibed in tap water for at least one hour, during which the mucilage of most species was able to fully expand (the average time to full expansion was 32 minutes).

**Table 1.** Meaning of symbols referenced in the study.

| **Parameter** | **Symbol** |
| --- | --- |
| Conversion factor (from Reynolds number to drag coefficient) | $\beta_1$ |
| Density of water | $\rho$ |
| Earth's gravity | $g$ |
| Max decayable mucilage mass | $m_{max}^*$ |
| Max mucilage mass | $m_{max}$ |
| Max projected seed area | $A$ |
| Mean projected area | $\bar{A}$ |
| Modified drag coefficient | $k_d$ |
| Mucilage decay coefficient | $D$ |
| Mucilage expansion speed coefficient | $a$ |
| Terminal velocity | $v$ |
| Time | $t$ |
| Total imbibed seed mass | $m_{total}$ |
| Total imbibed seed volume | $V_{total}$ |

**Figure 3.**
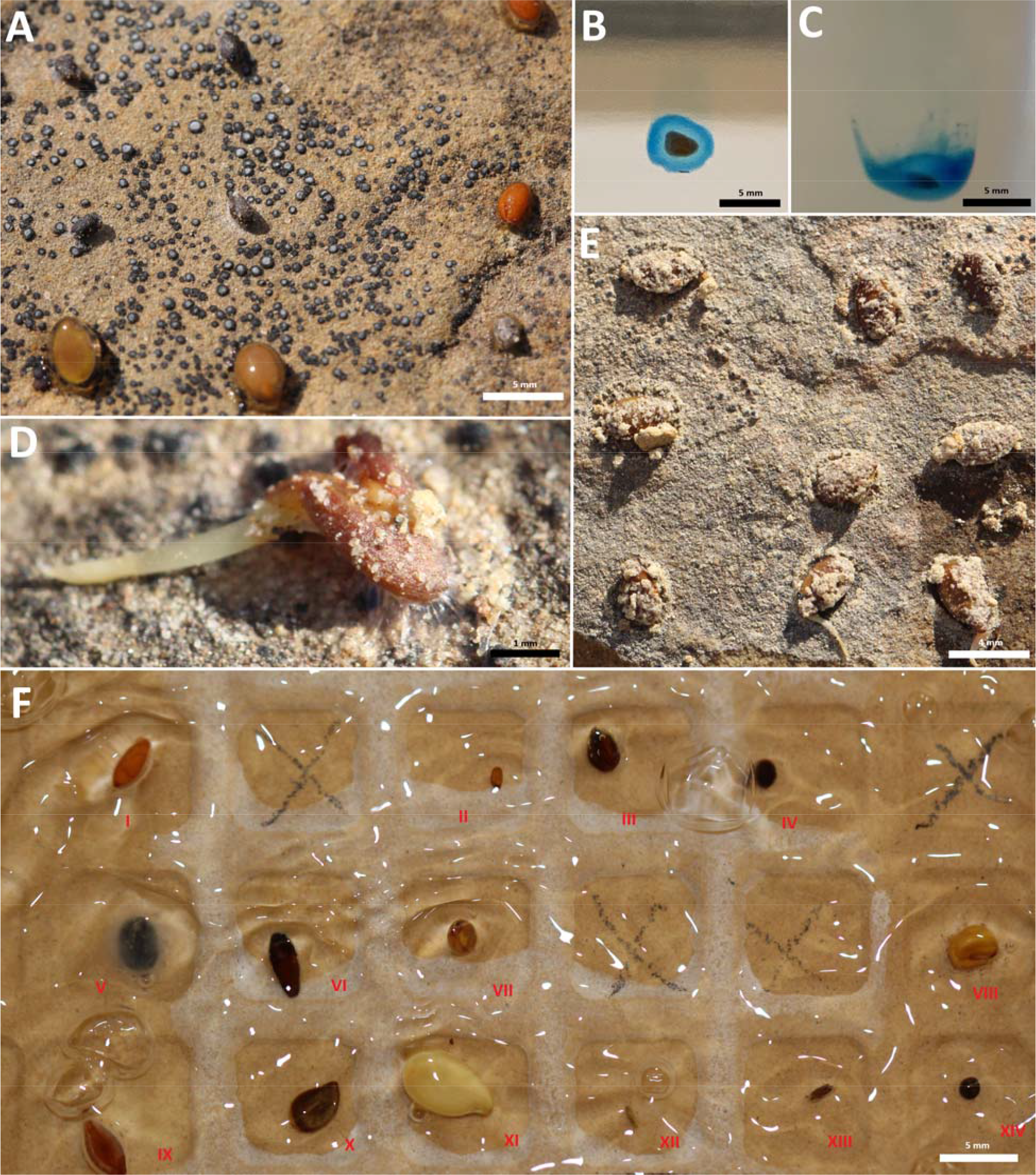
(A) Imbibed seeds with wet mucilage drying on sandstone (*Ocimum basilicum,* four in upper left; *Lepidium sativum*, two in upper right; *Linum grandiflorum,* two in lower left; *Salvia hispanica,* lower right). (B) *Gilia tricolor* and (C) *Dracocephalum* ‘blue dragon’ seeds stained with methylene blue falling in water. The mucilage envelope in C can be seen deformed and shearing off as it falls, in contrast to the limited deformation in B. (D) *Le. sativum* seed attached by dried mucilage strands on sandstone in the process of elongating its radical. (E) *Li. grandiflorum* with entrapped substrates attached to sandstone in the field, after a bout of natural surface flow. (F) Seeds attached to the back side of a tile during a dislodgement assay ((I) *Le. sativum;* (II) *Capsella bursa-pastoris;* (III) *Prunella grandiflora;* (IV) *Plectranthus scutellarioides*; (V) *O. basilicum;* (VI) *Plantago maritima*; (VII) *Eruca vesicaria;* (VIII) *Anastatica hierochuntica*; (IX) *Plantago ovata;* (X) *Linum perenne;* (XI) *Linum usitatissimum;* (XII) *Matricaria chamomilla;* (XIII) *Artemisia dracunculus;* (XIV) *Thymus vulgaris*).

Because previous wetting-drying cycles of loose seed bind them together and to objects, we can be reasonably sure that this was the first wetting-drying cycle for each seed in the experiment. We placed one imbibed seed of each species inside individual divots on the back side of each ceramic tile (10 × 10 cm Hudson Brilliant White Glossy Ceramic Wall Tile) in a predetermined randomized order and random orientation, then allowed the seeds to dry overnight (sufficient for complete drying of all species). We attached each tile in its predetermined orientation to a 10.5 × 10.5 × 12.5 cm plastic flowerpot with the bottom cut out, which acted as a funnel, and mounted on a 45° incline < 15 cm below individual water faucets. To present different conditions, alternating tiles were oriented right side up or upside down. To test a range of flow rates to get meaningful data from species with both weak and strong anchorage, each faucet was set to a flow rate between 6.7×10^-5^ - 1.4×10^-2^ L/s/cm. The flow from each faucet was measured at the beginning and end of each trial. As pressure changed based on other building users, the mean of the two values was recorded as the flow speed treatment (the small variations in flow rate over time within a faucet were dwarfed by the large variation between the set flow rates). The trials began when water started flowing and ended after either six or seven days. Each tile was monitored continuously for the first 15 minutes, on an hourly basis for the next five hours, and two to five times a day subsequently, during which we recorded the timing of each observed seed dislodgement. We used at most ten sinks simultaneously and ran the experiment over the span of two weeks for a total of 20 trials. One seed in one trial (*Leptosiphon ‘*hybrid’) was excluded due to a missing dislodgement time. An additional experiment that manipulated the drying time of seeds before exposure of erosive water flow is detailed in the supplement.

### Functional trait assays

To evaluate our hypothesized mechanisms of seed anchorage, we measured a variety of morphological and functional traits of the seeds. We measured dry seed mass for each species by weighing several seeds together and taking the mean for a single seed; this measurement was repeated twice per species. For each species, we also estimated the individual volume and mean projected area of four fully imbibed seeds for each species using a digital microscope (DinoCapture 2.0) by measuring the diameter of the three principal axes (see supplement). The dislodgement force of dry seeds was measured for 7-28 individual seeds/species using a modified method from Pan et al. (2021). We attached imbibed seeds to glass slides (United Scientific Supplies, model MSLF01) instead of adding water to dry seeds on petri dishes, which improved consistency and precision over our previous method. We also standardized the seed orientation, only pressing the Pesola spring scale perpendicularly against the long axis of the seed. Because the slipperiness of wet mucilage was not amenable to the same method, the dislodgement force of seeds when mucilage was wet was measured using a modified double-sided lever balance that pulled imbibed seeds apart from two pieces of filter paper (Whatman #1, 55mm, #1001-055, also see supplement; Figure S3). For this assay, we measured each species three times, each time using five seeds.

We characterized the quantity of mucilage produced by different species and the temporal dynamics of mucilage quantity in water. For short term dynamics, we measured the mass of imbibed seeds of each species at roughly 0.5, 2, 8, 32, 120, 480, and 1200 minutes after wetting. We weighed four seeds at each time point, discarding seeds after weighing as their mucilage structure may have been damaged during handling. For each species at each time point *t*, we found the mean species mucilage mass (*m*) by subtracting the mean species wet mass by the mean species dry seed mass. Then, using (1), modeled after Darcy’s law of hydrodynamic flow (supplemental method for derivation), we estimated mucilage mass of a fully imbibed seed (*m_max_*, which is functionally equivalent to mucilage mass after 1 hour for nearly all species) and the speed coefficient of mucilage secretion in s^-1^ (*a*):

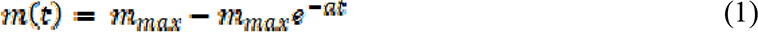

We repeated the assay a few times for species which had large posterior credible intervals around the estimated parameters.

To characterize the expanded mucilage decay over longer time periods when in water, we weighed four different seeds every day for seven days for each species. Twenty-eight seeds of each species were imbibed in 40 mL of tap water at 1 °C to which we added two drops of 1% bleach to prevent seed germination and decomposition (the cold water also slowed any bacterial growth, which is often rapid in submerged seeds). We estimated the rate of mucilage dissolution with (2):

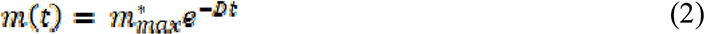

where *m(t)* is the mass of mucilage at time *t*, is the maximum mucilage mass that can decay, and *D* is the mucilage decay rate coefficient in days^-1^. We estimated as a free parameter and allowed it to differ from estimated from equation (1) as not all parts of mucilage may decay with time.

We further estimated a modified drag coefficient in water through a terminal velocity assay (Loudon & Zhang, 2002). In a 17 × 90 cm transparent acrylic cylinder filled to the brim with water at rest, we dropped imbibed seeds from the top and measured the time it took for each seed to sink to 60 cm depth. We repeated this assay ten times for each species. All seeds reached terminal velocity within the first ten centimeters of dropping distance, hence they were measured from 20 to 80 cm below the dropping point. From the equation for objects traveling at terminal velocity, we isolated an experimental constant (*k_d_*) that is in essence equivalent to the drag coefficient, but with the dependency on terminal velocity factored out (see supplement for derivation).

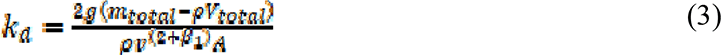

Here, *g* is the acceleration due to gravity, *m_total_* is the mass of the seed with mucilage, ρ is the density of water, *V_total_* is the volume of the imbibed seed with mucilage, β*_1_* is a conversion constant that relates the drag coefficient to Reynolds number, *A* is the projected area, and *v* is the terminal velocity of the seed. All parameters that we did not directly measure (*m_max_*, *a*, *D*, and *k_d_*) were estimated using individual species linear or non-linear models with moderately regularizing priors (see supplement).

### Components of seed anchorage

All continuous variables besides *D*, *k_d_*, seed mass, and time to dislodgement were log transformed to improve linearity and all continuous predictors were standardized as Z-scores. Unsupported interactions were dropped, and the models refitted. All statistical analyses were conducted using R (version 4.0.5; R Core Team 2020). All models were constructed using the Stan interface package *brms* (Bürkner, 2017, 2018) or *rstanarm* (Goodrich et al., 2020). All cross-validations were done using the package *loo* (Vehtari et al., 2017, 2019).

We first characterized the extent to which time to dislodgement was affected by a seed’s attachment to substrate and the degree of drag it experienced. We analyzed the time to dislodgement of each seed in a log-normal accelerated failure time (AFT) multilevel model (*n* = 1039 seeds). Tile orientation, flow speed, dislodgement force of wet mucilage, dislodgement force of dry mucilage, mean projected area, and the experimental drag constant *k_d_* were included as fixed effects. Flow speed was allowed to interact with the dislodgement force of wet and dry mucilage, mean projected area, and *k_d_* to test whether these processes have different effect sizes in different flow intensities. We also included mucilage decay *D* and mucilage wetting speed *a* as fixed effects and allowed them to interact with force and *k_d_* as the temporally variable quantity of mucilage might moderate the effect of force and *k_d_*. To take into account the uncertainty in estimated values of *k_d_*, *a,* and *D*, we performed multiple imputations at the species level using standard errors associated with each measurement. Individual tiles and species were included as random group intercepts. In order to account for some unmeasured variables shared among close relatives that affect the results, we performed a phylogenetic correction using a Brownian correlation structure. We constructed this matrix from a species level tree pruned from a synthesis of the most recent mega-phylogenies (Zanne, et al. 2014, Smith & Brown, 2018, *package*: *V.PhyloMaker* Jin & Qian, 2019). Unresolved species were assigned as polytomies at the genus basal node (Qian & Jin, 2016). *R^2^* was computed using equation three from Gelman et al. (2019) with or without group-level effects.

### Contributions of seed and mucilage traits to components of seed anchorage

To understand how seed and mucilage traits contribute to mechanistic component of fluid resistance and anchorage ability, we analyzed mean species dislodgement force of dry mucilage, dislodgement force of wet mucilage, mean projected area, and *k_d_* in four linear models as the response variable (*n* = 52). Dry seed mass and max mucilage mass *m_max_*were included in all models as fixed effects. Mucilage decay and an interaction with *m_max_* were included in the models for dislodgement force of wet mucilage and *k_d_*. Phylogenetic relationship was specified in all models as a correlation matrix. Measurement error multiple imputations were again conducted for variables we did not directly measure (*m_max_*, *a*, *D*, and *k_d_*).

### Structural equation model

We performed a piecewise confirmatory path analysis to characterize the relative magnitude and direction of how the studied seed functional traits mechanistically contributed to seed anchorage. Our structural equation model (SEM) was composed of five sub-models: four linear models for 1) dislodgement force of wet mucilage, 2) dislodgement force of dry mucilage, 3) mean projected area of mucilage envelope, and 4) *k_d_* that followed the same model structure as above, in addition to 5) an average species AFT model. We condensed individual seed level time to dislodgement into mean time to dislodgement for each species as the destructive nature of our trait assays necessitated that we only had mean species level trait data. Mean time to dislodgement for each species at mean flow speed was predicted from an AFT multilevel model as before, following the same model structure, but with only flow speed and tile orientation as fixed effects. Due to computational limitations, we limited multiple imputations to predicted dislodgement time and *k_d_*. Plausible alternative paths were checked by including each path in a new model and comparing the leave one out information criterion (*LOOIC*). Plausible alternative paths were rejected when the decrease in *LOOIC* was < 2.

### Marginal utility of functional assays

Finally, to determine what traits are most important to measure if time and resources are limited (for instance, in broad community functional trait studies), we characterized the marginal predictive value of different seed functional traits in understanding seed anchorage. Based on the results of our path analysis and regression models, we selected five seed traits (mean projected seed area, seed mass, *k_d_*, dislodgement force of wet and dry mucilage) for forward model selection. We began with an AFT multilevel model of individual seed dislodgement time as before but with only flow speed and tile orientation as fixed effects. Variables were selected based on grouped-13-fold cross-validation information criterion; each refit of the model left four species out of sample at a time (Roberts et al., 2017).

## Results

### Components of seed anchorage

We found strong evidence that dislodgement force and drag force were mechanistically driving the time to dislodgement (*Conditional R^2^* = 0.71, 95% *Credible interval* (*CI_95_*) = [0.68, 1.0], *Marginal R^2^* = 0.61, *CI_95_* = [0.53, 1.0]). Seeds on average remained on tiles for 5.3 days, although the time to dislodgement was highly variable among flow rates and among species. A standard deviation in the time to dislodgement spanned over two orders of magnitude. The ability for seeds to anchor to substrates substantially increased resistance to dislodgement under surface flow (Figure 4A-B). One *SD* increase in log dislodgement force of dry and wet mucilage was associated with a 2900% (β = 3.4 ± 0.74, *CI_95_* = [1.9, 4.8]) and 630% increase in time to dislodgement respectively (β = 1.8 ± 0.68, *CI_95_* = [0.50, 3.2]). Flow intensity did not affect the efficacy of either the dislodgement force of dry (β = -0.21 ± 0.26, *CI_95_* = [-0.74, 0.30]) or wet mucilage (β = 0.32 ± 0.27, *CI_95_* = [-0.21, 0.84]).

**Figure 4.**
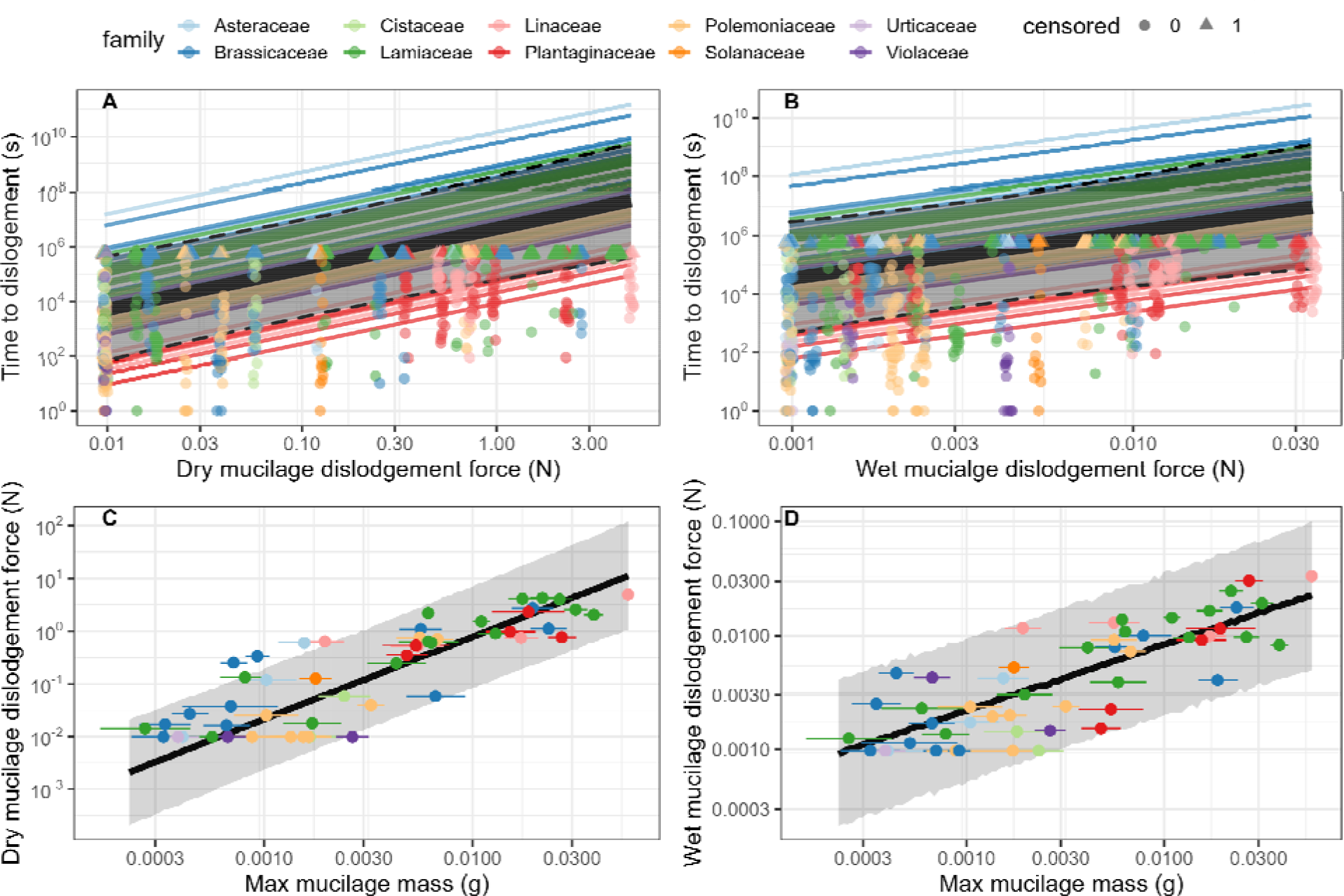
Relationship between time to dislodgement and dislodgement force (A & B) and between dislodgement force and fully imbibed mucilage mass (C & D) as predicted by a phylogenetic AFT multilevel model and two phylogenetic linear models (*n* = 1039). Black lines show the global relationship with grey ribbons representing the 95% *CIs*. Points and lines are colored by family. In A-B, species specific slopes are plotted as thinner lines. Each point corresponds to individual observed seed dislodgement time. Observation of dislodgement time is censored at one week; censored observations are indicated by triangles, whereas uncensored observations are indicated by circles. In C-D, each point corresponds to imputed measurement error free mean species mucilage mass and observed mean species dislodgement force. Error bars represent 95% *CIs*.

Variables that contributed to higher drag decreased time to dislodgement. The negative effect of the modified drag coefficient was strongest at lower flow intensities (β*_interaction_* = 0.41 ± 0.19, *CI_95_* = [0.057, 0.79]; Figure 5). At the 25^th^ and 75^th^ percentile flow speed, an *SD* higher *k_d_* was associated with an 84% and 44% reduction in time to dislodgement respectively (β*_kd_* = -1.2 ± 0.65, *CI_95_*= [-2.5, 0.084]). Flow speed had a negative effect on time to dislodgement regardless of *k_d_* (β*_flow_* = -1.8 ± 0.32, *CI_95_* = [-2.4, -1.1]). An *SD* higher flow speed reduced time to dislodgement by 83% when *k_d_* was held at average. Mean projected seed area did not have a significant effect on time to dislodgement (β = -0.57 ± 0.78, *CI_95_* = [-2.1, 0.96]) or interact with flow speed (β = -0.40 ± 0.27, *CI_95_* = [-0.92, 0.14]). In contrast to our hypotheses, neither mucilage decay (*D*) or wetting speed (*a*) affected time to dislodgement (β*_D_* = -0.55 ± 0.72, *CI_95D_* = [-1.9, 0.96]; β*_a_* = 0.49 ± 0.83, *CI_95a_*= [-1.2, 2.1]), or interacted significantly with the dislodgement force of wet mucilage (β*_D_* = 0.55 ± 1.1, *CI_95D_* = [-1.8, 2.4]; β*_a_* = -0.50 ± 0.93, *CI_95a_* = [-2.3, 1.4]), the dislodgement force of dry mucilage (β*_D_* = 0.054 ± 1.1, *CI_95D_* = [-2.2, 2.3]; β*_a_* = 0.44 ± 1.0, *CI_95a_* = [-1.5, 2.4]), or the drag coefficient, *k_d_* (β*_D_* = 0.82 ± 1.7, *CI_95D_* = [-3.9, 2.9]; β*_a_* = 0.077 ± 0.83, *CI_95a_* = [-1.6 1.6]).

**Figure 5.**
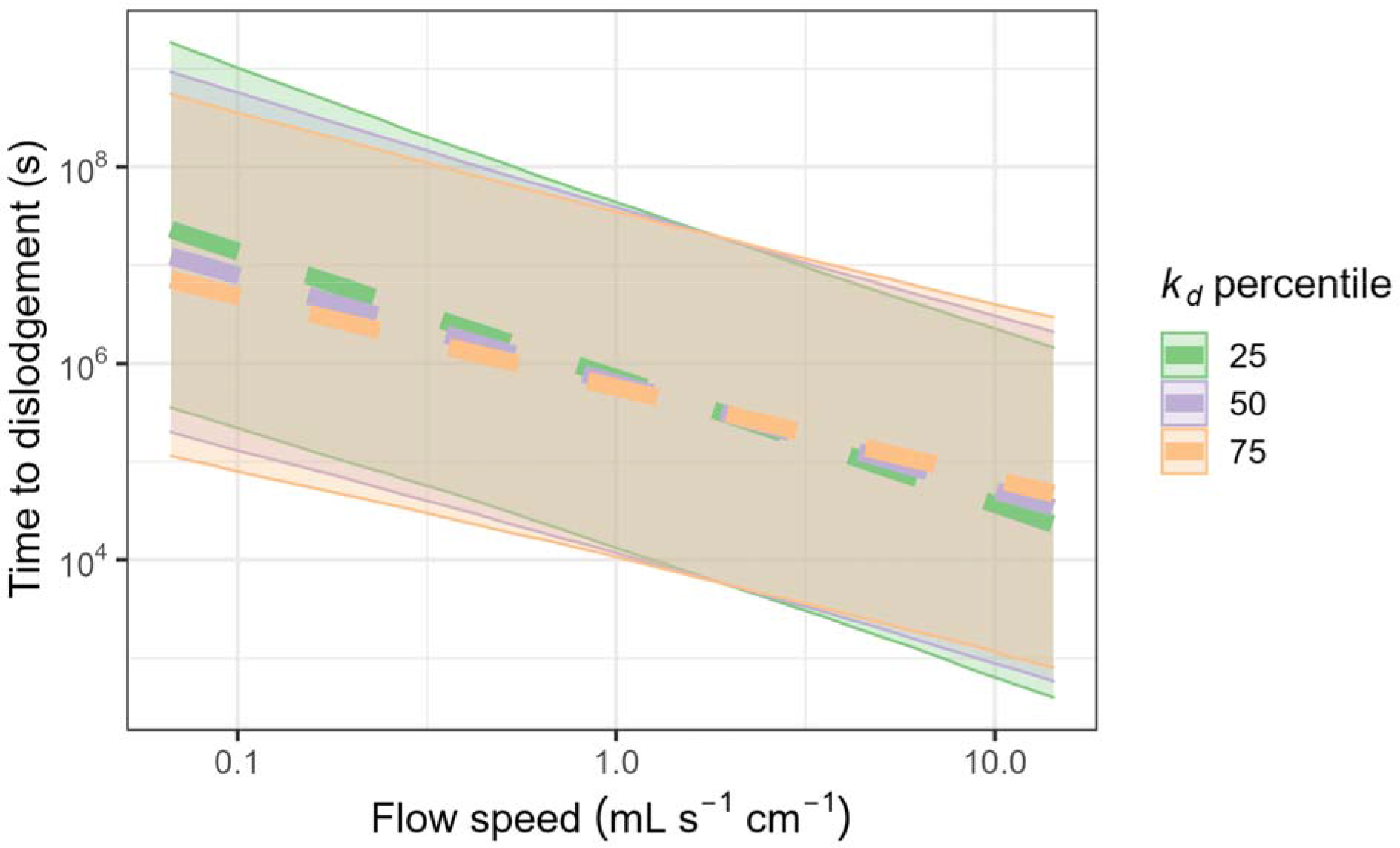
Effect of flow speed on time to dislodgement as predicted by a phylogenetic AFT multilevel model. Colors correspond to a seed with modified drag coefficient *k_d_* value at the 25^th^, 50^th^, or 75^th^ percentile. Dashed lines show the global trends with ribbons showing the 95% *CIs*.

Some seeds germinated during the trial. Dislodgement of the seedling post-germination occurred in 1% of seeds (11/1039) and a reanalysis, counting these germinated seeds as not dislodged, yielded results that were within rounding error from those presented here.

### Contributions of seed and mucilage traits to dislodgement force

Mucilaginous seeds anchored strongly to substrates, both after a wetting-drying cycle (i.e., when dry) and while mucilage was fully imbibed (i.e., when wet). Although the method of measurement was different, the dislodgement force greatly depended on the hydration status of the mucilage. The dislodgement force of mucilage to glass slides when dry averaged 0.13 N, while the dislodgement force of wet mucilage to filter paper averaged 0.0043 N. The amount of mucilage that seeds produced correlated well with the strength of substrate anchorage for both when the mucilage was dry (β = 2.3 ± 0.24, *CI_95_* = [1.9, 2.8]) and wet (β = 0.86 ± 0.13, *CI_95_* = [0.61, 1.1]) (Figure 4C-D). A one *SD* increase in log mucilage mass was associated with a 900% and 140% increase in the dislodgement force of dry and wet mucilage respectively. Higher seed mass was significantly correlated with lower dislodgement force of dry mucilage (β = -0.72 ± 0.30, *CI_95_* = [-1.3, -0.14]), but not with the dislodgement force of wet mucilage (β = -0.023 ± 0.15, *CI_95_* = [-0.33, 0.25]). Faster mucilage decay was positively correlated with higher dislodgement force of wet mucilage (β = 0.65 ± 0.17, *CI_95_* = [0.38, 1.0]) and its effect did not depend on the amount of mucilage (β = -0.17 ± 0.17, *CI_95_* = [-0.52, 0.15]). A one *SD* increase in the rate of mucilage decay corresponded to a 92% higher dislodgement force of wet mucilage. Together, these variables accounted for most of the variation in dislodgement force of dry mucilage (*R^2^*= 0.71, *CI_95_* = [0.63, 0.77]) and wet mucilage (*R^2^* = 0.76, *CI_95_* = [0.65, 0.85]).

### Contributions of seed and mucilage traits to drag coefficient and projected area

Mean projected area of imbibed seeds was significantly associated with higher amount of mucilage produced, independent of seed size. A one *SD* increase in log mucilage mass was associated with 90% higher mean projected area (β = 0.64 ± 0.068, *CI_95_* = [0.51, 0.77]). Seed mass was correlated positively with mean projected area, but the association was only marginally significant (β = 0.15 ± 0.086, *CI_95_* = [-0.020, 0.32]). Together, both variables explained most variation in mean projected area (*R^2^* = 0.89, *CI_95_* = [0.84, 0.92]). Our modified drag coefficient *k_d_* averaged 0.9, with some estimates extending into the negative domain as seed volumes were likely generally overestimated. Seed mass (β = 0.34 ± 0.49, *CI_95_* = [-0.60, 1.3]), mucilage mass (β = 0.19 ± 0.38, *CI_95_* = [-0.57, 0.94]), the rate of mucilage decay (β = 0.00070 ± 0.0078, *CI_95_* = [-0.015, 0.016]), and an interaction between the rate of mucilage decay and mucilage mass did not explain *k_d_* (β = -0.015± 0.015, *CI_95_* = [-0.043, 0.014]).

### Structural equation model

The best SEM is shown in Figure 6A. We found strong evidence that higher mucilage mass indirectly increased species’ antitelechorous potential through increasing dislodgement force of wet and dry mucilage, rather than directly affecting time to dislodgement (Δ*LOOIC* = 16). Through increasing both the dislodgement force of wet and dry mucilage, a one *SD* increase in log mucilage mass corresponded to a staggering 280-times increase in time to dislodgement, an effect which far overshadowed other seed traits in relative importance (Figure 6B). Both mucilage wetting speed and the rate of mucilage decay also had a positive indirect effect on time to dislodgement through increasing dislodgement force of wet mucilage, although the magnitude of their effect was comparatively minor (<10% the effect size of mucilage mass). Holding mucilage mass constant, seeds with a higher mass had a large negative indirect association with time to dislodgement through lower dislodgement force of mucilage. A one *SD* higher seed mass was associated with a 66% reduction in anchorage potential through lower dislodgement force of dry mucilage. The best model also included a direct path from seed mass to time to dislodgement (Δ*LOOIC* = 22), but the weak negative effect was not distinguishable from zero (β = -0.19 ± 0.15, *CI_95_*= [-0.49, 0.10]). At mean flow speed, the modified drag coefficient had a significant negative effect on time to dislodgement in the SEM, roughly equivalent in magnitude as the effect of the dislodgement force of wet mucilage. However, the modified drag coefficient was not affected by the amount of mucilage of the species. In contrast, while seed mucilage mass substantially increased mean projected seed area, mucilage mass ultimately did not indirectly reduce time to dislodgement through the mechanism of increased projected area because the effect of mean projected seed area is not distinguishable from zero.

**Figure 6.**
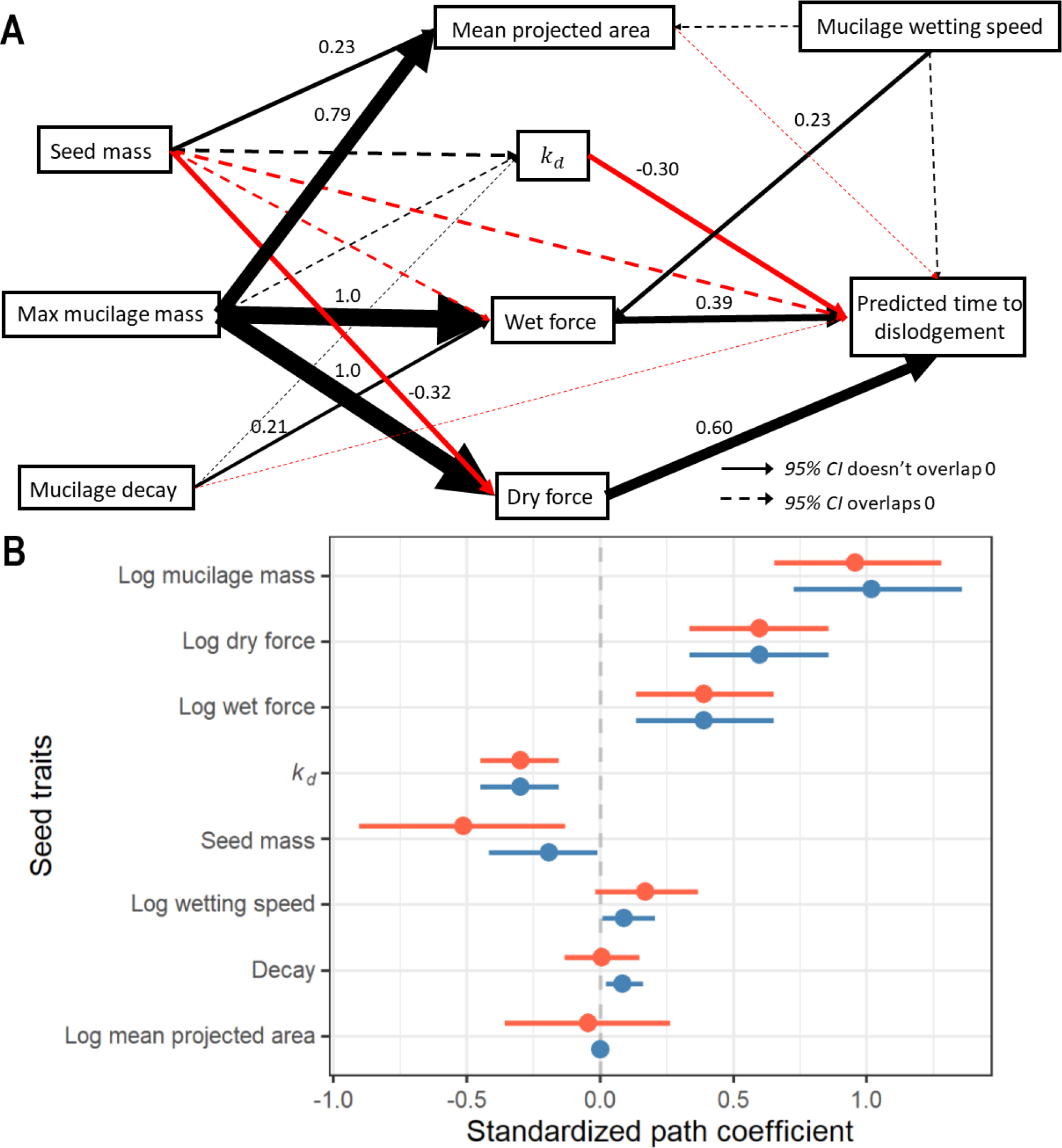
(A) Path diagram of the best SEM. Paths with *95% CI* overlapping with zero are indicated by dashed lines; otherwise, they are indicated by a solid line along with their corresponding standardized path coefficient. The size and color of all paths were scaled to the magnitude and direction of the effect respectively (red = negative; black = positive). A direct path from seed mass to dislodgement time (Δ*LOOIC* = 22), from wetting speed to mean projected area (Δ*LOOIC* = 6), and from wetting speed to dislodgement force of wet mucilage were added (Δ*LOOIC* = 23), though only the path from wetting speed to dislodgement force of wet mucilage was significant (β = 0.23 ± 0.10, *CI_95_* = [0.033, 0.43]). This positive association was likely because seeds that released mucilage faster had more mucilage at the time the dislodgement force was measured. (B) Total effect of each seed trait on predicted log species mean time to dislodgement. Error bars represent 95% *CIs*. Total path coefficients calculated based on all paths or only significant paths are shown in red and blue respectively. All variables besides mucilage decay rate, *k_d_*, and seed mass are on the log scale.

### Marginal utility of functional assays

Stepwise cross-validation revealed that including dislodgement force of dry mucilage was sufficient to achieve roughly three-quarters of the predictive performance of the best model containing dry dislodgement force, *k_d_*, dislodgement force of wet mucilage, and mean projected area (Figure S4). Including the next best variable *k_d_*brings the model’s predictive performance up to 93% of the best model.

## Discussion

Through explicitly testing our model of the physical mechanisms leading to dislodgement, we demonstrated that mucilage mass was by far the most important trait that affects mucilage-mediated seed anchorage and that it acts on time to dislodgement by increasing the force necessary for dislodgement. Thus, we suggest the dislodgement force assay of dry mucilage as a simple proxy for mucilage-mediated antitelechorous potential in future comparative studies of seed functional ecology (e.g. database such as Saatkamp et al., 2019), as this measurement had approximately 75% of the predictive performance of the best model and is very easily measured. Our mechanistic model, based on dislodgement force and fluid resistance, explained time to dislodgement well (*R^2^* = 0.71). Thus, this model (Figure 6A) is a useful heuristic to employ in further investigations of seed anchorage. We do not mean to suggest that other unmeasured variables (e.g. seed surface characteristics, mucilage composition, and mucilage structure) do not contribute meaningfully; rather, the basic measurements we performed are sufficient to capture the vast majority of the relative anchorage potential across a diversity of plants.

### What drives dislodgement force?

Dislodgement force, especially that of dried mucilage was the most important predictor of dislodgement resistance (Dried mucilage dislodgement: partial *R^2^* = 0.26, *CI_95_* = [0.00012, 0.50], Wet mucilage dislodgement: partial *R^2^* = 0.076, *CI_95_*= [-0.094, 0.31]). It correlated strongly with mucilage mass, a result we also found with a different set of species and a slightly different dislodgement assay (Pan et al., 2021), suggesting that increased dislodgement force can be driven by simply increasing mucilage production. Mucilage mass may be quite variable among close relatives (Table S2). A particularly striking example occurred within *Dracocephalum* (Lamiaceae). An ornamental variety of *D. moldavica* had 39 mg of mucilage in a large envelope around the nutlet and a dislodgement force of > 2 N, while the similarly sized nutlets of *D. parviflorum* had < 1 mg of mucilage localized in small projections at one end, and an unmeasurably small dislodgement force (scored as 0.0098 N in this analysis).

However, considering mucilage mass as the primary predictor is overly simplistic; other factors, probably composition and seed shape, are likely to contribute to the dislodgement force of dry mucilage. The pectic mucilage of *Linum* spp., rich in rhamnogalacturonan and arabinoxylan with unusual side group substitutions, provides a good example (Naran et al., 2008). Dislodgement force of the wet and dry mucilage of Linaceae was consistently underpredicted by mucilage mass, suggesting that this mucilage composition may have a stronger dislodgement force than other types (also suggested by Kreitschitz et al., 2021a). In contrast, seeds of species of Polemoniaceae were more easily dislodged than expected given their mucilage mass. Their mucilage composition and primary function of mucilage in nature is thus far unknown but deserves more investigation. We have found that it provides a defensive benefit against harvester ants to several Polemoniaceae species (LoPresti et al., 2019, Pan et al., 2021), though it was similarly not as protective compared to other families in those studies.

The substrate that seeds attach to is also important. We measured dried mucilage dislodgement force on glass microscope slides for maximum replicability. However, when testing substrates for this experiment, we found that seeds that were attached to increasingly finer grit sandpaper dislodged with less force (unpublished data). Fortunately, the glass slide assay was a good proxy for the performance on the tiles, as evidenced by its high explanatory power, although there could be realistic substrates where it is far less applicable. Drying speed also alters dislodgement force and mucilage structure (personal observations), though it is unclear how this interacts with substrate, and therefore deserving of increased scrutiny. How mucilage mass, seed shape, and mucilage composition change dislodgement force across substrate types and environmental conditions are entirely unexplored questions that could increase ecological realism in future studies of mucilage-mediated antitelechory.

### Why was drag a less important predictor of dislodgement?

The drag coefficient, despite having a significant effect, was a relatively poor predictor of time to dislodgement in our study (partial *R^2^* = 0.085, *CI_95_*= [-0.044, 0.27]). There are both theoretical and methodological explanations for this result. It is likely that for most seeds in the most realistic erosive conditions (i.e. not our highest flow rates), drag is not sufficient to overcome the strong anchorage to the substrate (i.e. there may be a threshold effect at which point drag matters). Our estimates, derived from seeds falling at terminal velocity in a water-filled cylinder, also do not properly reflect the flow regime or mucilage envelope dynamics during erosive conditions. The mucilage envelope probably deforms to a much higher extent in higher flow rates and outer layers may shear off (Figure 3B-C), thereby reducing drag below our estimates. Further, in higher and more turbulent flow regimes, theory predicts the drag coefficient rapidly decays (White, 1991), thus contributing a lesser extent to the overall drag. For those reasons, our estimates of this coefficient were probably inaccurate, though we believe that they reasonably reflect relative species differences in this study. As such, that the effect of the modified drag coefficient was strongest at lower flow rates was hardly surprising. Deeper investigation into this matter is beyond the scope of this study and more complex drag dynamics are surely at play under field conditions than were considered here. For instance, in field pilot trials, the attached seeds caught substrate and plant debris during times of surface flow which surely affected the seeds’ fluid resistance (Figure 3D-E). Finally, the poor explanatory performance of mucilage envelope mean projected area might be attributed to similar causes. In addition, we noticed that water rarely flowed over the top of larger seeds except in higher surface flow conditions (Figure 3F). Thus, only a fraction of the estimated projected area was truly interacting with water.

### Why were mucilage decay and expansion rate poor predictors of dislodgement?

Given that the attachment potential of wet and dry mucilage differed so greatly, and that mucilage mass was an important predictor of attachment potential (Pan et al., 2021), we hypothesized that the time the mucilage envelope takes to expand and the rate of decay when submerged would be important predictors of time to dislodgement. Each of these metrics varied greatly between species (Table S2). The mucilage of *Prunella grandiflora* (Lamiaceae) decayed far more quickly (18%/day) than any other species (mean: 5%/day), and some species had no measurable decay during the week-long trial. Time to 95% mucilage expansion varied greatly as well, from just three minutes in *Lobularia maritima* (Brassicaceae) to 5.5 hours in *Linum grandiflorum* (Linaceae). Even with these large differences, both traits were poor predictors of time to dislodgement (partial *R^2^* = 0.042, *CI*_95a_ = [-0.064, 0.18]; partial *R^2^* = 0.043, *CI* = [- 0.058, 0.18]).

We hypothesized that mucilage decay in water would reduce the dislodgement force of wet mucilage and drag, chipping away the mucilage anchorage over time. However, the rate of mucilage decay appeared to be generally sufficiently low as to be of little importance within the span of one week. Though, because seeds often need to withstand erosive forces sporadically over months in nature (during which time the mucilage may be fed on by microbes or small invertebrates: Buse & Filser, 2014) before an adequate growing condition arrives, the compounding mucilage decay could take on a greater importance over longer time scales and in natural settings. In other cases, the presence of an adherent layer that is possibly more decay resistant in the mucilage envelope may continue to provide sufficient anchorage even after the non-adherent layer is gone (Zhao et al., 2017).

Similarly, we hypothesized that mucilage expansion rate would affect antitelechorous potential by moderating the switch between the much stronger dislodgement force of dry mucilage and the much weaker dislodgement force of wet mucilage. However, our results suggest that the attachment provided by either dry or wet mucilage was likely sufficient to anchor most species against most erosive surface flow in nature. In an additional experiment, seeds with wet mucilage that were not allowed time to dry remained equally resistant to dislodgement as seeds that were and the length of mucilage drying time did not explain time to dislodgement (see supplement). Considering that mucilage expansion speed may increase the dislodgement force of wet mucilage (Figure 6A), its possible role in facilitating more rapid onset of mucilage-mediated seed anchorage, even in situations of limited moisture exposure time (e.g. morning dew), may be critical in nature.

### Potential adaptive significance of seed mucilage

Across plants, particularly from desert or ruderal environments, extremely limited dispersal is a common phenomenon, which likely decreases seed loss to unsuitable habitats (Ellner & Schmida, 1981). Cheplick (2022) termed this limited dispersal from mother plants “philomatry” and differentiates extrinsic (mostly environmental) and intrinsic (mostly plant characters) drivers. Anchorage by seed mucilage prevents dislodgement (Garcia-Fayos et al., 2013, Engelbrecht et al., 2014), and therefore can be considered an intrinsic driver of philomatry. Dispersal, or anti-dispersal, traits are often part of broader syndromes correlated with other life- history traits across plants (Cheplick, 2022). The presence and amount of mucilage has broader correlates, including ruderal life-histories (Grubert, 1974, Ryding, 2001), and being more pronounced in warmer areas with high solar radiation (Pan et al., 2021). We hypothesize that this trait may allow exploitation of an erosive niche where pre-germination seed anchorage may be advantageous. In addition, seed mucilage may help seeds avoid becoming badly abraded in the erosion process (e.g. with a coating in LoPresti et al., 2019) or avoid deeper burial with lower chance of emergence (Mennan & Zandstra, 2006). In environments where other plants are continuously removed by erosion, the remaining plants may face reduced competition, compared to areas where seeds collect after erosion in high numbers. A lower plant density also likely reduces disease pressure (Bell et al., 2006) and herbivory (Kim & Underwood, 2015). Thus, anchorage mechanisms could be seen as essential traits for certain plants to uniquely exploit highly erosive niches – though the broader correlations may underlie other evolutionary drivers.

It should be noted, however, that seeds need to anchor using the mucilage layer only until they germinate and begin root growth. Thus, time to dislodgement (*t_D_*), as well as traits that contribute to it, for a given seed can only be understood ultimately in light of its time to germination (*t_G_*), with success defined as:

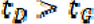

Accordingly, for species that germinate quickly, lower anchorage ability need not be an indictment of its ability to colonize highly erosive environments or the efficacy of the species’ anchorage mechanisms. Recent meta-analysis, however, revealed that seeds with limited dispersal generally have delayed germination time (Chen et al. 2020). Some authors have also reported that the presence of seed mucilage delayed germination (e.g. Khajeh-Hosseini & Mostashar-Shahidi, 2016, Souza et al., 2020), further suggesting mucilage mediated antitelechory may be a broader syndrome and seed anchorage is an important component of the particular life history strategy.

Anchorage, though probably an important function of mucilage in nature (Garcia-Fayos et al., 2013, Engelbrecht et al., 2014), also cannot be understood in isolation, as seed mucilage is a highly multi-functional trait. Some of the characteristics that we measured here also contribute to other beneficial effects of mucilage (e.g. mucilage mass correlates with germination under water stress conditions: Teixeira et al., 2020; dislodgement force of dry mucilage correlates with resistance to granivory: Pan et al., 2021). Additionally, it is reasonable, though as of yet untested, that these same traits contribute more to other functions: dislodgement force of wet mucilage probably drives epizoochorous dispersal (Ryding, 2001), a slower mucilage decay rate may increase survival of seeds through the gut of herbivores (Kreitschitz et al., 2021a), and faster imbibition rate could lead to better DNA repair during brief wetting periods (Huang et al., 2008). These all may thus be part of a syndrome of life-history traits that species with mucilaginous seeds largely share and possibly account for their similar ecological niches (Grubert, 1974), a result that may suggest a mucilaginous life-history syndrome, of which limited dispersal is just one component (à la Cheplick, 2022).

To understand the drivers of mucilage evolution would require a more comprehensive sampling of species and more extensive measurement of traits that uniquely account for the many alternative ecological functions. Increasingly finer mechanistic understanding of the seed and mucilage traits that predict specific ecological functions would create more opportunities for teasing apart the evolutionary drivers of those specific traits, the other life-history correlates, and seed mucilage as a broader trait itself. Such an effort would be necessary for causal inference of the selection pressures responsible for the evolution of mucilage on seeds.

## Supporting information

Supplemental methods and results

## Acknowledgements

The USDA ARS GRIN program provided some of the seeds and were very prompt and helpful in coordination and shipment; special thanks to Jeffrey Carstens, Tiffany Fields, and Alex Sanchez. The computing for this project was performed at the High-Performance Computing Center at Oklahoma State University supported in part through the National Science Foundation grant OAC-1531128. We thank Jesse Schafer, head of operations, who was tremendously helpful in speeding up our script. We also thank Bill Henley and Andrew Doust for sharing their lab space for us to run our experiment, John Pan for providing technical guidance on fluid mechanics, the Wetzel lab and Rick Karban for providing manuscript feedback, and Mark Fishbein for both sharing sinks and commenting on our manuscript. VSP is supported by the Early Start fellowship from Michigan State University. EFL is supported by startup funds from Oklahoma State University.

## Author contributions

VSP, CG, & EFL designed the study, collected the data, and made the figures. VSP conducted the analysis. VSP & EFL wrote the manuscript. CG provided editorial support.

## Data availability statement

All data and code used to generate this manuscript are available on Dryad and Zenodo respectively: https://doi.org/10.25338/B81M00

