## Supplemental methods and results for "Anchorage by seed mucilage prevents seed dislodgement in high surface flow: a mechanistic investigation"

*Seed anchorage pilot experiment*

To understand whether mucilage facilitates ground anchorage in highly erosive environments, we initially conducted a field pilot experiment with the seeds of *Lepidium sativum*, *Linum grandiflorum*, *Salvia hispanica*, and *Ocimum basilicum*. On March 9, 2021, we placed 25 seeds of each species in 5 × 5 grids next to each other for a total of 16 blocks on an outcrop of slightly incline sandstone at McPherson Reserve in Stillwater, Oklahoma (degree decimals: 36.100997, -97.205267). The seeds in half of the blocks were placed on the sandstone without imbibition (control), the other half were imbibed for >1 hour to release the mucilage (treatment). We monitored whether the seeds stayed in place at least once a week for a month (until the start of spring in April). Time to dislodgement was analyzed with a log-normal accelerated failure time multilevel model (*n* = 1600). Treatment was included as a predictor and allowed to vary by species. Species and blocks were included as random intercept terms.

*Wet dislodgement force measurement*

We developed a simple low-cost method for measuring the dislodgement force of wet mucilage, suitable for relative measurement across a wide range of species. The double-sided lever balance is composed of a bar resting on a pivot with a 5-dram and 40-dram vial suspended with string on either end (Figure S3). To the bottom of the 5-dram vial and the top of a glass slide, we each tied a piece of filter paper (Whatman grade 1; Whatman International Ltd, Springfield Mill James Whatman Way Kent, ME14 2LE United Kingdom) that acts as a standardized surface upon which the wet mucilage adheres to. However, one might easily as well replace the filter paper with another substrate to test ecologically relevant situations for wet dislodgement (e.g. bird feathers or mammal fur for epizoochory hypotheses).

To measure the dislodgement force of we mucilage, we pressed five seeds, arranged in a ring each 1 cm apart, between the two pieces of filter paper firmly, binding them together. On the other side, we added water into the 40-dram vial slowly until the dislodgement failed. We then removed any seed still stuck to the 5-dram vial and blow dried the filter paper. Next, we added water to the empty 5-dram vial until the vial returned to its original position before failure (because of friction in the pivot, we also pipetted excess water out once the vial returned to position). The minimum weight of water in the 5-dram vial that kept the vial in its original position was then measured with a balance. We divided the weight of the water by the number of seeds used to obtain a mean dislodgement force of wet mucilage per seed. This mean has the caveat that it may be slightly higher than if each seed were to be measured individually; seeds do not get dislodged as a group in the field. A minimum of 0.5 g (0.0049 N) was recorded for each water weight measurement. The filter paper was blow dried or replaced after every use. The entire instrument can be made with readily available supplies. We built ours out of an old file organizer and some tape, but any lever arm that rotates easily on a fulcrum should work just fine.

*Seed volume and projected area*

For simplicity, we assumed that all seeds are shaped as an ellipsoid. We estimated the volume of a fully imbibed seed *V_total_* with

$V_{total}=\frac{\pi}{6}L_{total}W_{total}T_{total}$

(4)

where *L, W*, and *T* are the three perpendicular diameters of the seed with mucilage in the order of size. The slipperiness of mucilage was not amenable to direct measurement of the thickness, thus, *T_total_* was estimated with

$T_{total}=W_{mucilage}+\left( T_{DrySeed}\times\frac{W_{WetSeed}}{W_{DrySeed}} \right)$

(5)

where we add the mean species mucilage width to the thickness of the dry seed multiplied by some seed expansion factor that accounts for the seed absorbing water. The mean projected area was estimated using the formula

$\bar{A}_{P}=\frac{S}{4}$

(6)

where *S* is the surface area approximated by

$S=4\pi{(\frac{\left( \frac{L}{2} \right)^{p}\left( \frac{W}{2} \right)^{p}+\left( \frac{L}{2} \right)^{p}\left( \frac{T}{2} \right)^{p}+{(\frac{T}{2})}^{p}{(\frac{W}{2})}^{p}}{3})}^{\frac{1}{p}}$

(7)

where *p* is a constant equal to 1.6075.

*Describing mucilage quantity with time: expansion and decay*

Mucilage is released by bursting of mucilage secreting cells upon water absorption (North et al., 2014) or by rehydration of dried mucilage. In any case, we can expect that the net rate of water movement into the mucilage matrix is regulated by the water potential of mucilage relative to that of the environment. This dynamic takes the form of Darcy’s law of hydraulic flow (8), which has a history of being used to model seed water absorption (Vertucci, 1989).

$$Q=-K\left( \psi_{seed}-\psi_{medium} \right)$$

(8)

Where *Q* is the flow rate of water, *K* is the hydraulic conductivity coefficient (diffusive permeability factor), and *ψ* is the water potential. The conservation of mass suggests that *Q* must be proportional to the rate of change of the mucilage mass. Then, assuming that *ψ* is proportional to the mucilage quantity, we can rewrite (8) to describe the rate of change of mucilage mass

$\frac{dm}{dt}=-a\left( m-m_{max} \right)$

(9)

where *m* is the instantaneous mass of mucilage, *m_max_* is the maximum amount of mucilage when *ψ_mucilage_ = ψ_water_,* and *a* is a factor that describes how easily water is absorbed by the mucilage or mucilage secreting cells, related mostly to mucilage composition, cell wall permeability, and surface area. Integrating by time *t* and solving for *m(t = 0) = 0*, we obtain an equation for mucilage mass at any given time:

$$m\left( t \right)= m_{max}-m_{max}e^{-at}$$

(1)

This model fits our data to a satisfactory degree. Our median model has a mean absolute percentage error of 24%, and most of the error came from the intraspecific variability in seed and mucilage mass. However, the model is only suitable for short-term characterization of mucilage mass within a day, as mucilage may begin to break down under extended imbibition (Naran et al., 2008). Most seeds we tested retained most of their mucilage integrity for a few days, even months. For the process of mucilage decay, we assume that the rate of decay is proportional to the amount of mucilage *m* that is breaking down and some decay constant *D* that describes how quickly some amount of mucilage breaks down:

$\frac{dm}{dt}=-Dm$

(10)

Integrating by *t* and assuming $m\left( t=0 \right)=m_{max}^{*}$ as mucilage expansion is no longer relevant at this timescale, we obtain an equation that describes the mucilage mass over the long term:

$m\left( t \right)= {m_{\max}^{*}e}^{-Dt}$

(2)

Out of simplicity, equation (2) assumes that the amount of mucilage that can decay decays over time at a constant rate. However, some species have both an adherent and non-adherent mucilage layer (Zhao et al., 2017), where the former is likely more resistant to decay. The mucilage decay coefficient should therefore be interpreted as a time averaged index of mucilage loss.

*Deriving the modified drag coefficient*

Our goal in this section is to estimate a single propensity index for drag for each seed after accounting for projected seed area. Because the dominating mechanistic factors in drag depend on the flow regime, we begin by calculating the Reynolds number for seeds in our terminal velocity assay. We use (11)

$Re=\frac{\rho vL_{total}}{\mu}$

(11)

where *Re* is the Reynolds number, *ρ* is the density of water (997 kg/m^3^), *µ* is the dynamic viscosity of water (8.9×10^-4^ kg/m/s), *v* is the velocity of the seed, and *L_total_* is the longest diameter of the seed including the fully expanded mucilage. The *Re* of our species fell within the intermediate range of 6 to 120. At this flow regime, the force of drag (*F_D_*) can be approximated by (12)

$F_{D}=\frac{1}{2}{\rho C}_{d}Av^{2}$

(12)

where *ρ* is the density of water, *A* is the largest projected area of the seed as per our observation of the seed orientation, *v* is the seed velocity, and *C_d_* is the drag coefficient. Unfortunately, at an intermediate *Re* range, *C_d_* cannot be treated as a constant. It is dependent on *Re*, which is in turn dependent on the seed velocity. There are numerous approximations of *C_d_* even with many simplifying assumptions (Beetstra et al., 2007, Barry & Parlange, 2018). We will use (13) supplied by White (1991; 3-225), as it is a simple approximation that has precedence in the ecological literature (Loudon & Zhang, 2002).

|  | $C_{d}=\frac{24}{Re}+\frac{6}{1+\sqrt{Re}}+0.4$; for $0 < Re \leq{2 \times10}^{5}$ | (13) |
| --- | --- | --- |

Our goal is to estimate a single propensity index for drag without the influence of irrelevant assay specific variables. However, as one can see from (13), the velocity term inside *Re* cannot be isolated and factored out. We must remove the effect of terminal velocity as it is a result of many interacting forces, which are irrelevant to drag propensity and different for each species. In the seed anchorage experiment, *v* would be approximately the flow speed, not the terminal velocity. Fortunately, for the range of *Re* we are working with, *Re* and *C_d_* have a strong linear relationship on the log scale (Figure S5). We thus assume the relationship:

|  | $ln\left( C_{d} \right) \approx\beta_{1}ln\left( Re \right) +\beta_{0}$; for $6 \leq Re \leq120$ | (14) |
| --- | --- | --- |

where *β_0_* and *β_1_* are conversion constants that relate the drag coefficient to Reynolds number. Simulating 10,000 points using (13) revealed that with *β_0_ ≈* 2.752 and *β_1_ ≈* -0.5738, (14) can approximate the result of (13) within an 8.4% mean absolute percent error. This is good enough for our purposes as measurement error and intraspecific variability are at least one magnitude higher. Thus, armed with (11) and (14), we can rewrite (12) in a form that is easier to work with:

|  | $F_{D}\approx\frac{1}{2}\rho e^{\beta_{0}}\left( \frac{\rho vL_{total}}{\mu} \right)^{\beta_{1}}Av^{2}$; for $6 \leq Re \leq120$ | (15) |
| --- | --- | --- |

Returning to our seed dropping assay, we describe the forces acting on a seed falling in water at terminal velocity with

$$F_{net}=0=F_{g}-F_{B}-F_{D}$$

(16)

where *F_g_* is the gravitational force, *F_B_* is the buoyant force, and *F_D_* is the drag force. We substitute with known formulae

$0=m_{total}g-\rho gV_{total}-\frac{1}{2}\rho e^{\beta_{0}}\left( \frac{\rho vL_{total}}{\mu} \right)^{\beta_{1}}Av^{2}$

(17)

where *g* is the acceleration due to gravity (9.8 m/s^2^), *m_total_* is the mass of the seed with mucilage, *V_total_* is the volume of the imbibed seed with mucilage.

Removing the irrelevant variables from (17), what is left is defined as our modified drag coefficient *k_d_*

$k_{d}=\frac{2g\left( m_{total}-\rho V_{total} \right)}{\rho v^{\left( 2+\beta_{1} \right)}A}$

(3)

Based on (17), one can find that $k_{d}\approx e^{\beta_{0}}\left( \frac{\rho L_{total}}{\mu} \right)^{\beta_{1}}$, although only approximately, as the influence of many other factors contained within the experimental constant *k_d_*, such as skin friction and mucilage viscosity, are not explicitly modeled in (17).

We used different measured *m_total_* values for *Nicandra physalodes* in this assay compared to the others as the batch of seeds used in the dropping assay were more mucilaginous than the ones used in other assays.

*Extracting coefficients from data*

We fitted our mathematical models to the data of each species individually to extract species specific experimental coefficients. Equation (1) was fitted as a non-linear regression model of the form

$$m_{i}=m_{max}-m_{max}\exp\left( at_{i} \right)+\epsilon_{i}$$

(18)

where $m_{i}$ is the *i*th measurement of mucilage mass, $m_{max}$ is the estimated max mucilage mass, $a$

is the mucilage expansion speed coefficient, $t_{i}$ is the imbibition time of the *i*th measurement, and $\epsilon_{i}$ is the error of the *i*th measurement.

Equation (2) was fitted as a linear regression model after log transforming the mucilage mass.

$$\ln\left( m_{i} \right)=\beta_{intercept}+\beta_{slope}day_{i}+\epsilon_{i}$$

(19)

Here, $\beta_{intercept}$ is equivalent to $\ln\left( m_{max}^{*} \right)$, $\beta_{slope}$ is equivalent to $-D$, $m_{i}$ is the *i*th measurement of mucilage mass, $day_{i}$ is the day of the *i*th measurement, and $\epsilon_{i}$ is the error of the *i*th measurement.

Equation (3) was fitted as a linear regression model that is forced through the origin after some reparameterization.

$$V_{i}^{'}=\beta_{slope}X_{i}+\epsilon_{i}$$

(20)

Here, $V_{i}^{'}$ is equivalent to $V_{total}-\frac{m_{total}}{\rho}$ , $X_{i}$ is equivalent to $\frac{-A{v_{i}}^{2+\beta_{1}}}{2g}$, $\beta_{slope}$ is equivalent to $k_{d}$, and $\epsilon_{i}$ is the error of the *i*th measurement. We parameterized the model this way for a few reasons. First, a linear model is easier to fit by Stan’s MCMC sampler. Second, it allows us to impose a specific regularizing prior on $\beta_{slope}$. Most importantly, $V_{i}^{'}$ is approximately normally distributed and measured with the most error, so separating it out allows us to model its error specifically.

*Drying time experiment*

It is possible that in some situations, mucilaginous seeds might not be able to dry for weeks during wet weather; in contrast in deserts, some species may dry extremely quickly. Considering the dislodgement potential of wet mucilage is magnitudes lower than when dry, it raises the questions whether the period of drying time is important for the resistance of seeds to dislodgement. We thus repeated two trials of our laboratory seed anchorage assay with five species of moderate propensity to dislodgement (i.e. the species does not immediately dislodge, nor does it never dislodge over a period of seven days: *Le. sativum*, *Li. grandiflorum*, *O.* *tenuiflorum*, *Plantago ovata*, *S. hispanica*). Instead of allowing seeds to become fully dry, for each species, we allowed the seeds to dry for 6 to 151 minutes at roughly 15-minute intervals (*n* = 10/species) in the laboratory (~22 C, 50% RH). Time to dislodgement was again analyzed with a log-normal AFT multilevel model (*n* = 100). Drying time was included as a fixed effect and allowed to vary by species. Tile and species were included as random intercepts. Phylogenetic relatedness was accounted for with a Brownian correlation matrix.

**Supplemental Result and Discussion**

*Seed anchorage pilot experiment*

Anchorage to the ground via mucilage was very effective against erosive forces in the field. Seeds that were imbibed and allowed to stick to sandstone remained in place on average for 32 days, roughly ten times longer than the non-imbibed seeds (*β* = 2.3 ± 0.49, 95% *CI* = [1.1, 3.2]). While some of the time to dislodgement differences between imbibed and non-imbibed seeds might be in part driven different susceptibility to seed predation (Pan et al., 2021), we did not observe any seed predation taking placing during the first hour of the experiment when the vast majority of seeds without mucilage were displaced by wind. In other pilot experiments, we were consistently surprised by the resilience of these mucilaginous seeds against dislodgement in other highly erosive environments, including a vertical clay cliff and a frequently flooded riverbank.

*Drying time experiment*

We found no evidence that longer drying time contributed to resistance to dislodgement (*β* = 0.42 ± 0.53, 95% *CI* = [-0.71, 1.5]). Thus, it appears that minimal drying is sufficient for seed mucilage to anchor seed to substrate, preventing dislodgement.

**Table S1.** Sources of seeds.

| **Family** | **Species** | **Source** |
| --- | --- | --- |
| Asteraceae | *Artemisia dracunculus* | Outsidepride |
|  | *Cladanthus arabicus* | Outsidepride |
|  | *Matricaria chamomilla* | Sustainable organic seeds, Trade wind fruits |
| Brassicaceae | *Anastatica hierochuntica* | Commercial |
|  | *Capsella bursa-pastoris* | Seeds Of Strength |
|  | *Cardamine hirsuta* | McPherson Reserve, Stillwater, Oklahoma (collected April 2021) |
|  | *Diplotaxis tenuifolia* | Hazzard’s seeds |
|  | *Eruca vesicaria* | Trade winds fruits |
|  | *Heliophila longifolia* | Hazzard’s seeds |
|  | *Ionopsidium acaule* | Hazzard’s seeds |
|  | *Lepidium campestre* | PI 633248 |
|  | *Lepidium meyenii* | Trade winds fruits |
|  | *Lepidium sativum* | Hemani |
|  | *Lobularia maritima* | Wildseed Farms |
| Cistaceae | *Helianthemum nummularium* | Outsidepride |
|  | *Helianthemum variable* | PI 292853 |
| Lamiaceae | *Dracocephalum 'blue dragon'* | Hazzard’s seeds |
|  | *Dracocephalum parviflorum* | W6 44097, W6 43742 |
|  | *Melissa officinalis* | Trade winds fruits |
|  | *Ocimum americanum* | Trade winds fruits |
|  | *Ocimum basilicum* | Sustainable Seed Company |
|  | *Ocimum tenuiflorum* | Botanical interests |
|  | *Plectranthus scutellarioides* | Seedman |
|  | *Prunella grandiflora* | Outsidepride |
|  | *Salvia coccinea* | Wildseed Farms |
|  | *Salvia columbariae* | Field collection, McLaughlin Reserve, Lake County, California |
|  | *Salvia farinacea* | The Harvest of Jonny Wildseed |
|  | *Salvia hispanica* | Better body foods organic chia seeds |
|  | *Salvia rosmarinus* | Hazzard’s seeds, Outsidepride |
|  | *Salvia sclarea* | Outsidepride |
|  | *Thymus vulgaris* | Botanical Interests |
| Linaceae | *Linum grandiflorum* | Sustainable Seed Company |
|  | *Linum lewisii* | Sustainable Seed Company |
|  | *Linum perenne* | Botanical Interests |
|  | *Linum usitatissimum* | Premium Gold Whole flax seed |
| Plantaginaceae | *Plantago arenaria* | W6 4760 |
|  | *Plantago erecta* | McLaughlin Reserve, Lake County, California (collected June 2018) |
|  | *Plantago lanceolata* | McLaughlin Reserve, Lake County, California (collected June 2018) |
|  | *Plantago maritima* | PI 415825 |
|  | *Plantago ovata* | Seedman |
| Polemoniaceae | *Gilia leptantha* | Davis, California (collected June 2018) |
|  | *Gilia tricolor* | Outsidepride |
|  | *Leptosiphon 'hybrida'* | Outsidepride |
|  | *Linanthus grandiflorus* | Outsidepride |
|  | *Polemonium caeruleum* | Hazzard’s seeds |
|  | *Polemonium pauciflorum* | Hazzard’s seeds |
|  | *Polemonium viscosum* | Hazzard’s seeds |
|  | *Polemonium yezoense* | Hazzard’s seeds |
| Solanaceae | *Nicandra physalodes* | Hazzard’s seeds |
| Urticaceae | *Urtica dioica* | W6 33123 |
| Violaceae | *Viola tricolor* | Trade winds fruits, Botanical Interests |
|  | *Viola x wittrockiana* | Outsidepride |

**Table S2**. Measured and estimated mean species seed traits. (DM) Dry mass in mg; (TIM) total imbibed volume in mm^3^; (PA) mean projected area in mm^2^; (DF) dry mucilage dislodgement force in mN; (WF) wet mucilage dislodgement force in mN; (MM) max mucilage mass per seed in mg; (ESC) mucilage envelope expansion speed coefficient in s^-1^; (TE) time to 95% mucilage envelope expansion in minutes; (D) mucilage decay coefficient in day^-1^; (MDC) modified drag coefficient in (s/m)^β_1^; (DIS) proportion of seeds dislodged after one week.

| **Species** | **DM** | **TIV** | **PA** | **DF** | **WF** | **MM** | **ESC** | **TE** | **D** | **MDC** | **DIS** |
| --- | --- | --- | --- | --- | --- | --- | --- | --- | --- | --- | --- |
| *Artemisia dracunculus* | 0.20 | 0.85 | 1.1 | 9.8 | 0.98 | 0.41 | 0.0036 | 14 | 0.0036 | -1.4 | 0.55 |
| *Cladanthus arabicus* | 0.34 | 1.4 | 1.7 | 600 | 4.2 | 1.5 | 0.00092 | 55 | 0.017 | 1.4 | 0.0 |
| *Matricaria chamomilla* | 0.094 | 1.2 | 1.4 | 12 | 1.7 | 1.1 | 0.0051 | 9.7 | 0.10 | -0.77 | 0.15 |
| *Anastatica hierochuntica* | 2.0 | 21 | 9.7 | 1100 | 18 | 23 | 0.00055 | 91 | 0.057 | 1.6 | 0.0 |
| *Capsella bursa-pastoris* | 0.096 | 0.74 | 1.0 | 27 | 4.7 | 0.43 | 0.0047 | 11 | 0.053 | -2.3 | 0.0 |
| *Cardamine hirsuta* | 0.11 | 0.83 | 1.2 | 17 | 2.5 | 0.33 | 0.0030 | 17 | 0.033 | -3 | 0.20 |
| *Diplotaxis tenuifolia* | 0.24 | 0.79 | 1.1 | 9.8 | 0.98 | 0.31 | 0.0017 | 30 | 0.0099 | -0.9 | 0.90 |
| *Eruca vesicaria* | 1.7 | 6.4 | 4.2 | 1100 | 7.9 | 5.6 | 0.0024 | 21 | 0.056 | 0.61 | 0.0 |
| *Heliophila longifolia* | 0.34 | 0.92 | 1.5 | 250 | 0.98 | 0.70 | 0.0015 | 32 | 0.010 | 0.24 | 0.80 |
| *Ionopsidium acaule* | 0.34 | 2.3 | 2.1 | 16 | 1.7 | 0.65 | 0.015 | 3.4 | 0.038 | -2.7 | 0.95 |
| *Lepidium campestre* | 1.7 | 9.5 | 5.4 | 58 | 10 | 7.6 | 0.0025 | 20 | 0.046 | -0.061 | 0.40 |
| *Lepidium meyenii* | 0.46 | 0.71 | 1.1 | 37 | 1.1 | 0.52 | 0.0033 | 15 | 0.077 | 0.74 | 0.85 |
| *Lepidium sativum* | 2.2 | 21 | 9.4 | 2700 | 4.1 | 19 | 0.0021 | 24 | 0.015 | 0.21 | 0.25 |
| *Lobularia maritima* | 0.33 | 0.94 | 1.4 | 330 | 0.98 | 0.92 | 0.017 | 3.0 | 0.022 | 1.0 | 0.15 |
| *Helianthemum nummularium* | 1.2 | 6.0 | 4.0 | 58 | 0.98 | 2.5 | 0.0039 | 13 | 0.094 | -0.77 | 0.90 |
| *Helianthemum variable* | 1.3 | 2.5 | 2.3 | 9.8 | 1.4 | 1.8 | 0.0018 | 29 | -0.039 | 0.29 | 0.95 |
| *Dracocephalum* 'blue dragon' | 2.0 | 35 | 13 | 2000 | 8.3 | 38 | 0.0013 | 38 | 0.023 | 2.7 | 0.050 |
| *Dracocephalum parviflorum* | 1.5 | 3.2 | 2.9 | 9.8 | 2.3 | 0.55 | 0.0033 | 15 | 0.076 | -0.55 | 0.65 |
| *Melissa officinalis* | 0.57 | 1.8 | 1.9 | 18 | 3 | 2.1 | 0.0056 | 8.8 | 0.067 | 2.4 | 1.0 |
| *Ocimum americanum* | 1.5 | 25 | 10 | 4300 | 25 | 21 | 0.0036 | 14 | 0.035 | -1.1 | 0.0 |
| *Ocimum basilicum* | 0.76 | 24 | 10 | 4100 | 16 | 17 | 0.0032 | 16 | 0.074 | -1.9 | 0.05 |
| *Ocimum tenuiflorum* | 0.64 | 8.9 | 5.2 | 2200 | 14 | 6.1 | 0.0075 | 6.7 | 0.076 | -2.8 | 0.0 |
| *Plectranthus scutellarioides* | 0.28 | 0.81 | 1.1 | 14 | 1.3 | 0.23 | 0.0055 | 9.1 | 0.091 | -1.5 | 0.55 |
| *Prunella grandiflora* | 1.8 | 8.9 | 5.3 | 250 | 7.9 | 4.3 | 0.0020 | 25 | 0.20 | -1.8 | 0.25 |
| *Salvia coccinea* | 1.4 | 23 | 9.8 | 4000 | 9.7 | 26 | 0.0013 | 37 | 0.038 | 2.9 | 0.050 |
| *Salvia columbariae* | 0.9 | 7.0 | 4.6 | 600 | 11 | 6.3 | 0.0019 | 26 | 0.053 | 0.79 | 0.0 |
| *Salvia farinacea* | 1.5 | 6.5 | 4.3 | 600 | 3.9 | 6.1 | 0.0039 | 13 | 0.077 | 1.5 | 0.15 |
| *Salvia hispanica* | 1.0 | 13 | 6.8 | 900 | 9.6 | 13 | 0.0019 | 27 | -0.0053 | 1.2 | 0.25 |
| *Salvia rosmarinus* | 1.6 | 16 | 7.9 | 1500 | 14 | 11 | 0.0028 | 18 | 0.075 | -2.3 | 0.10 |
| *Salvia sclarea* | 3.8 | 36 | 13 | 2500 | 19 | 31 | 0.0030 | 16 | 0.0099 | -0.29 | 0.050 |
| *Thymus vulgaris* | 0.24 | 1.5 | 1.6 | 130 | 1.4 | 0.78 | 0.0046 | 11 | 0.034 | -2.2 | 0.25 |
| *Linum grandiflorum* | 3.2 | 23 | 11 | 4900 | 33 | 56 | 0.00015 | 330 | 0.096 | 6.7 | 0.80 |
| *Linum lewisii* | 1.6 | 2.8 | 3.7 | 500 | 13 | 5.5 | 0.0051 | 9.8 | 0.073 | 3.0 | 0.80 |
| *Linum perenne* | 1.4 | 2.8 | 3.5 | 640 | 12 | 1.9 | 0.0079 | 6.3 | 0.10 | 0.37 | 0.95 |
| *Linum usitatissimum* | 5.8 | 23 | 11 | 740 | 9.9 | 17 | 0.00095 | 52 | 0.085 | 0.053 | 0.85 |
| *Plantago arenaria* | 1.7 | 25 | 11 | 970 | 9.1 | 16 | 0.00082 | 61 | 0.082 | -1.5 | 0.85 |
| *Plantago erecta* | 1.6 | 27 | 11 | 760 | 30 | 27 | 0.00046 | 110 | 0.062 | 0.5 | 0.65 |
| *Plantago lanceolata* | 1.3 | 7.4 | 4.8 | 530 | 2.3 | 5.3 | 0.0032 | 16 | 0.022 | -0.41 | 0.85 |
| *Plantago maritima* | 0.80 | 5.6 | 4.0 | 350 | 1.5 | 4.9 | 0.0072 | 6.9 | 0.12 | 0.050 | 0.80 |
| *Plantago ovata* | 1.9 | 15 | 7.7 | 2300 | 12 | 19 | 0.0013 | 39 | 0.020 | 2.4 | 0.90 |
| *Gilia leptantha* | 1.1 | 15 | 7.6 | 700 | 7.3 | 6.8 | 0.0020 | 25 | 0.087 | -2.9 | 0.20 |
| *Gilia tricolor* | 0.50 | 6.8 | 4.4 | 760 | 9.2 | 5.4 | 0.0044 | 11 | 0.095 | -1.4 | 0.050 |
| *Leptosiphon* 'hybrida’ | 0.29 | 1.6 | 1.8 | 9.8 | 0.98 | 0.96 | 0.0011 | 44 | -0.00085 | -0.70 | 0.84 |
| *Linanthus grandiflorus* | 0.50 | 4.0 | 3.2 | 25 | 2.4 | 1.0 | 0.0041 | 12 | 0.063 | -2.5 | 0.60 |
| *Polemonium caeruleum* | 0.89 | 4.2 | 3.4 | 9.8 | 0.98 | 1.7 | 0.0012 | 42 | 0.011 | -1.2 | 0.90 |
| *Polemonium pauciflorum* | 1.0 | 4.8 | 3.6 | 39 | 2.4 | 3.2 | 0.00060 | 84 | 0.13 | -0.37 | 0.85 |
| *Polemonium viscosum* | 1.3 | 3.7 | 3.3 | 9.8 | 2.0 | 1.6 | 0.0023 | 22 | -0.034 | -0.46 | 0.95 |
| *Polemonium yezoense* | 0.95 | 2.3 | 2.4 | 9.8 | 2.0 | 1.4 | 0.0024 | 21 | 0.074 | 0.0068 | 0.90 |
| *Nicandra physalodes* | 0.70 | 11 | 5.6 | 130 | 5.3 | 1.7 | 0.0075 | 6.7 | 0.091 | 2.6 | 0.55 |
| *Urtica dioica* | 0.20 | 0.34 | 0.7 | 9.8 | 0.98 | 0.38 | 0.0098 | 5.1 | -0.0039 | 2.0 | 1.0 |
| *Viola tricolor* | 0.56 | 0.69 | 0.98 | 9.8 | 4.3 | 0.65 | 0.0044 | 11 | 0.033 | 1.9 | 0.85 |
| *Viola x wittrockiana* | 1.1 | 5.1 | 3.6 | 9.8 | 1.5 | 2.7 | 0.00056 | 89 | -0.008 | -1.4 | 0.65 |

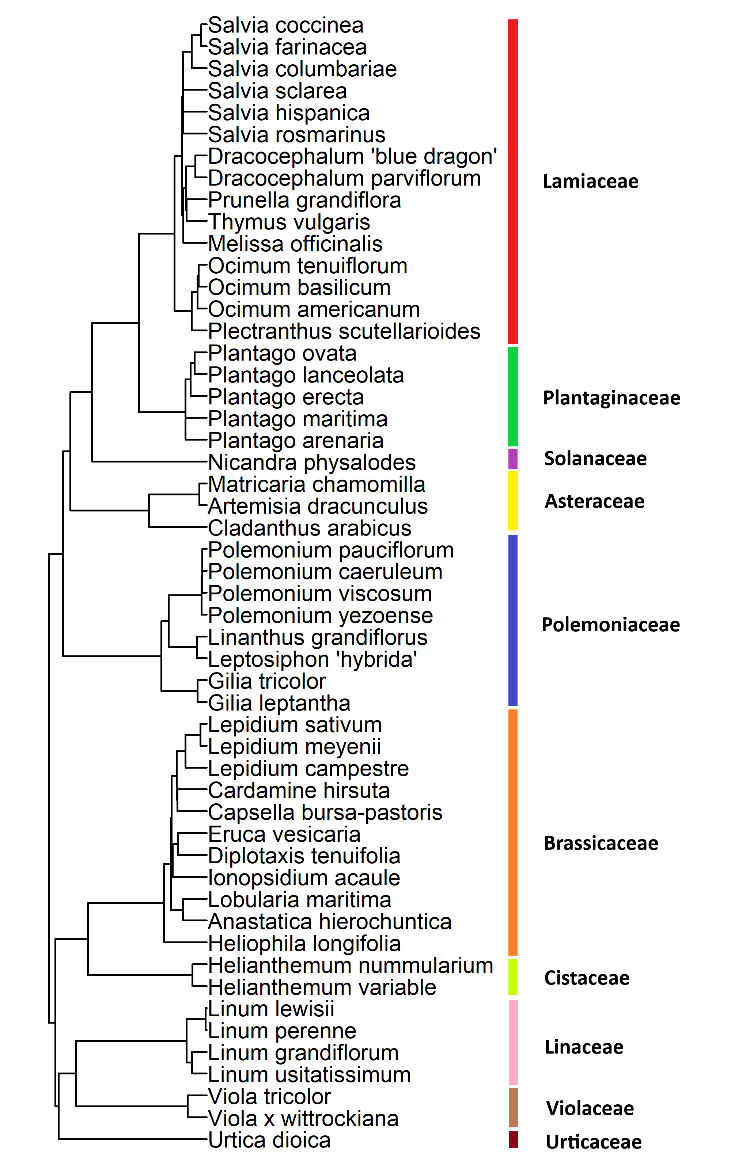

**Figure S1**. Species level phylogeny of the 52 species tested in this study. Unresolved placements are treated as polytomies at the genus basal node.

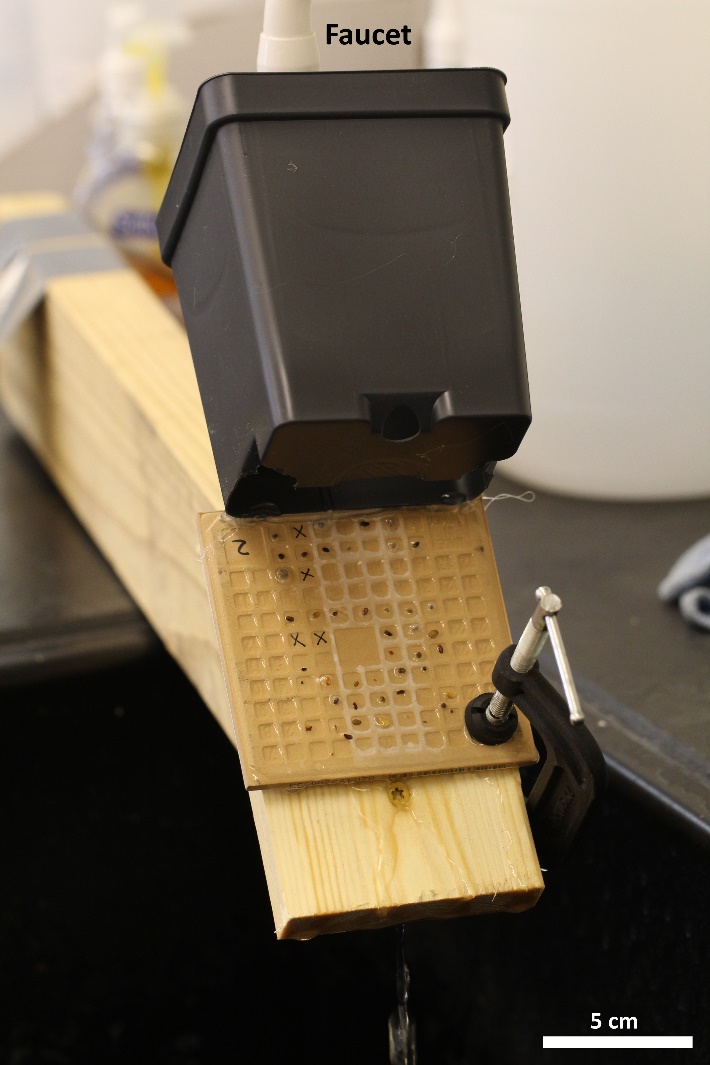

**Figure S2**. Seed anchorage assay setup. This particular tile was mounted upside down. We placed seeds in the visible squares on the back side of the tile (divots).

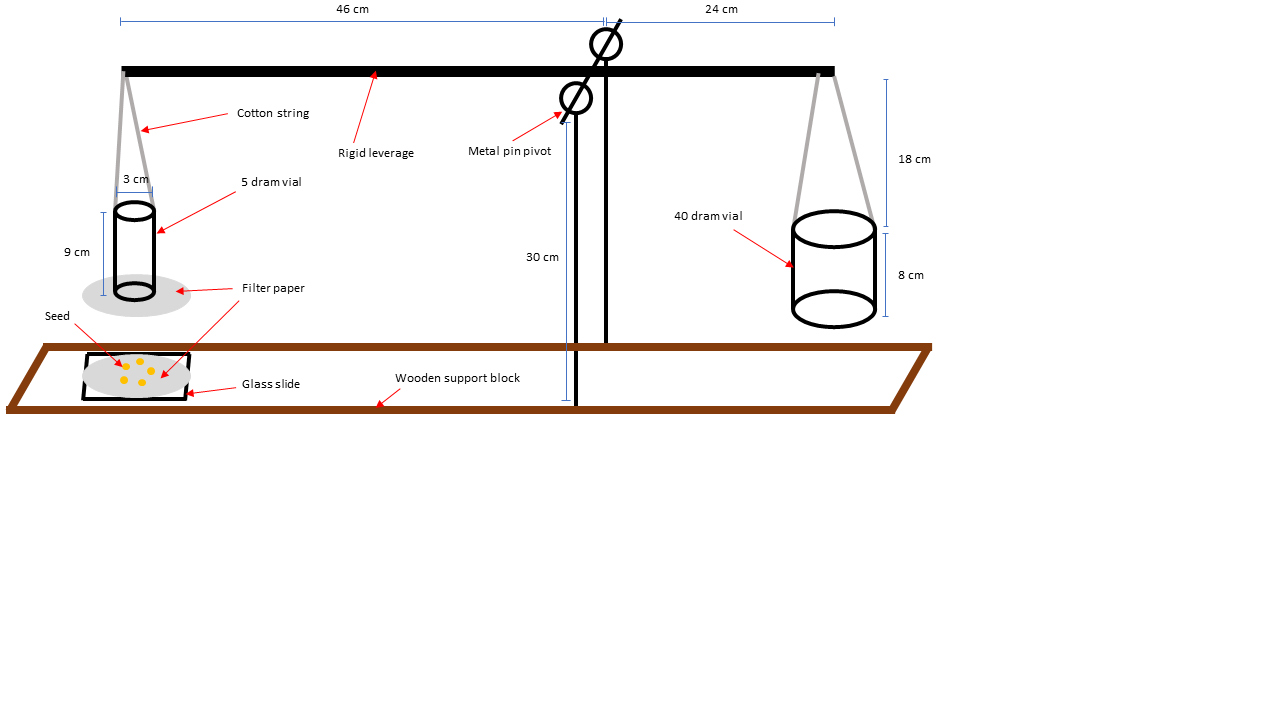

**Figure S3.** Modified double sided lever balance used to measure the dislodgement force of wet mucilage.

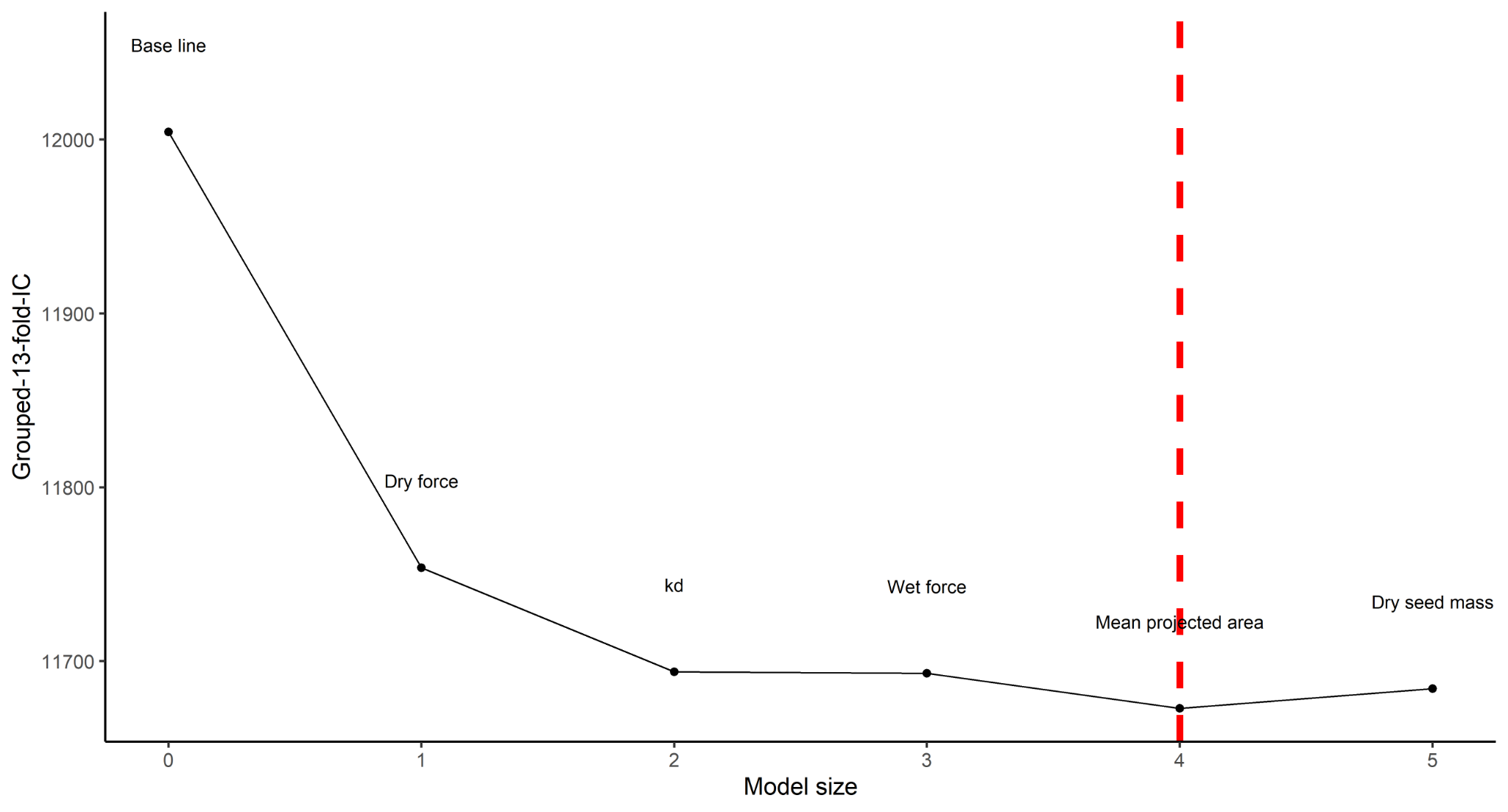

**Figure S4**. Model predictive performance in forward selection. Each new variables added are shown above the line. The baseline model contains log flow speed and tile orientation as fixed effects, and species and tile as random intercepts. Best model is indicated by the red dash line. The grouped-13-fold-IC is related to expected log pointwise predictive density (ELPD) by a factor of -2 and can thus be interpreted as a measure of model predictive performance.

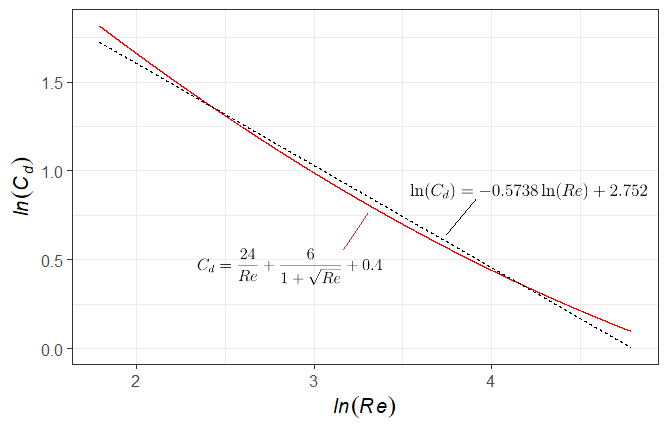

**Figure S5.** Relationship between the drag coefficient and Reynolds’ number for 6 ≤ *Re* ≤120. Red line is based on equation 3-225 from White (1991), which should be within 10% accuracy for a sphere in laminar flow with 0 < *Re* ≤ 2 × 10^5^. Dotted line is the rough approximation of the relationship based on (8).
